# Low pathogenic avian influenza virus infection retards colon microbiome diversification in two different chicken lines

**DOI:** 10.1101/2021.03.15.435422

**Authors:** Klaudia Chrzastek, Joy Leng, Mohammad Khalid Zakaria, Dagmara Bialy, Roberto La Ragione, Holly Shelton

## Abstract

A commensal microbiome regulates and is in turn regulated by viruses during host infection which can influence virus infectivity. In this study, analysis of colon microbiome population changes following a low pathogenicity avian influenza virus (AIV) of the H9N2 subtype infection of two different chicken breeds was conducted. Using 16S rRNA sequencing and subsequent data analysis we found reduced microbiome alpha diversity in the acute period of AIV infection (day 2-3) in both Rhode Island Red and VALO chicken lines. From day 4 post infection a gradual increase in diversity of the colon microbiome was observed, but the diversity did not reach the same level as in uninfected chickens by day 10 post infection, suggesting that AIV infection retards the natural accumulation of colon microbiome diversity, which may further influence chicken health following recovery from infection. Beta diversity analysis indicated differences in diversity between the chicken lines during and following acute influenza infection suggesting the impact of host gut microflora dysbiosis following H9N2 influenza virus infection could differ for different breeds.

## Introduction

Avian influenza A viruses (AIV), belong to the Orthomyxoviridae family, have segmented, single-stranded, negative sense RNA genomes with enveloped virions [1]. Based on pathogenicity, AIV can be categorised as low and high pathogenicity AIVs. Among low pathogenicity AIV (LPAIV), the H9 subtype circulates globally in wild birds and is endemic in domestic poultry in many countries in the Middle Eastern, Africa and Asia [2–8]. The majority, approximately 75%, of natural H9 isolates, are paired with the N2 neuraminidase (NA) subtype and are most frequently isolated from chickens, followed by waterfowl, pigeons, quail, and turkeys [9]. Infected chicken flocks usually experience mild respiratory distress, diarrhoea, decreased body weight in broilers, and a drop in egg production in layer hen flocks, the mortality rates are generally low, below 20% [10–14]. However, infected poultry are more susceptible to secondary infections, including bacterial infection and in such cases the mortality rate can increase up to 65% [15–18]. It has been shown that certain bacteria, such as *Staphylococcus* spp. can enhance influenza virus activation by indirect proteolytic cleavage and activation of the hemagglutinin (HA) using the virulence factor, staphylokinase [19].

The interplay between pathogens and host microbiome play an important role in health and disease in many vertebrates [25–28]. Compelling evidence has shown that the gut microbiome can play a role in pathogenesis of various human diseases including those with primary involvement outside of the gut, such as respiratory, renal, or neurologic [20–22]. For instance, recent studies reveal that immune protection and severity of infection by gammaherpesvirus, which can cause severe vasculitis and lethal pneumonia or respiratory syncytial virus infection of the lungs, can be dependent on the profile of the human gut microbiome [23, 24].

Studies in the chicken model have shown compositional changes in gut microbiome and differences in some of immune gene expression levels following H9N2 avian influenza virus (AIV), Newcastle disease virus (NDV) or infectious bronchitis virus (IBV) infection [29–32]. However, there are many environmental factors, such as age, breed, diet, housing, hygiene, and temperature that can also affect chicken microbiome [33, 34] and thus might change the interactions between the host and viruses during the time of infection. Furthermore, a recent murine study, demonstrated how polymorphism in host genes shape the intestinal microbiome and how host genetics influence the output microbiome by comparing genetically identical and genetically diverse mouse models [35]. It has been shown in chicken model that genetic background can influence viral pathogenesis [36–40]. For instance, two different inbred chicken lines, Fayoumi and Leghorn, have been used to evaluate mechanisms of genetic response to several different pathogens. [41, 42]. Deist, et al. [38] showed that Fayoumis chickens infected with NDV had a faster viral clearance than Leghorns chickens and higher serum antibody levels.

There are many factors contributing to the complex interplay between pathogens and host microbiome. Understanding of changes in host microbiome resulting from viral infections is particularly of interest, as it could provide information for developing new methods for infectious disease prevention and treatment. In this study we used two different chicken breeds, Rhode Island Red and VALO leghorns, reared in different facilities, to assess colon microbiome changes following H9N2 AIV infection. We analysed the commonality and differences in the host microbiome changes during and following infection of these two chicken lines.

## Results

### H9N2 AIV infection of chickens causes mild clinical signs

In both of our experiments, all birds survived H9N2 AIV challenge. RIR and VALO H9N2 infected chickens showed mild lethargy and diarrhoea especially between day 2 and day 4 post-challenge. There were no significant differences in body weight between control and infected birds for either breed (Supplementary Figure. 1). There were significant differences in body weight between the RIR and VALO chickens. At day 0 the RIR control group were on average 54.31 g (±5.5 g SEM) heavier than the VALO control group (p <0.0001) whereas at day 14 the RIR control group was 180.3 g (±38.2 g SEM) heavier than the VALO control group (p = 0.0421) (Supplementary Figure 1).

### VALO chickens shed H9N2 virus infection from the buccal cavity, a day longer than RIR chickens

Buccal viral shedding was determined by testing oropharyngeal (OP) swabs at day two, day three, day four, day five and day six post-challenge by plaque assay on MDCK cells (Figure 1). In experiment 1 where only RIR chickens were infected, all chickens shed virus from the OP cavity on day two with the average titre shed being 6.2×10^4^ pfu/ ml (±12730 pfu/ ml SEM). Virus titre declined on day four and day five, with no virus being recovered from samples taken on day six post infection (Figure 1A). In experiment 2, both RIR and VALO chickens were challenged with the same dose of the same H9N2 virus. At day two post challenge, all chickens in both lines shed virus from the OP cavity with no statistical difference in titre shed between the chicken lines being observed (average titres being 4.3 ×10^3^ pfu/ ml (± 921 pfu/ ml SEM) for RIR and 2.7 ×10^3^ pfu/ ml (±1086 pfu/ ml SEM) for VALO). Similarly, to experiment 1 virus titres in the OP cavity declined on days four and five with no observed virus shedding on day six post challenge for either line. We did see differences in the rate of viral clearance between the RIR and VALO chicken lines from day four post challenge onwards. On day four the mean OP virus shed was 1.6 ×10^3^ pfu/ ml (± 1129 pfu/ ml SEM) for the RIR compared to 3.6 ×10^3^ pfu/ ml (± 1121 pfu/ ml SEM) for the VALO (p value = 0.2465). On day five only 1 out of 8 RIR birds shed virus to titres above the limit of detection for the plaque assay whereas 7 out of 8 VALO birds did (average virus shed was 1.3 ×10^2^ pfu/ ml) (Figure 1B). Cloacal viral shedding of virus by all the birds in both experiments was conducted by qRT-PCR for viral M gene, but only a single RIR bird on day four post challenge had a positive result (data not shown).

**Figure 1.**
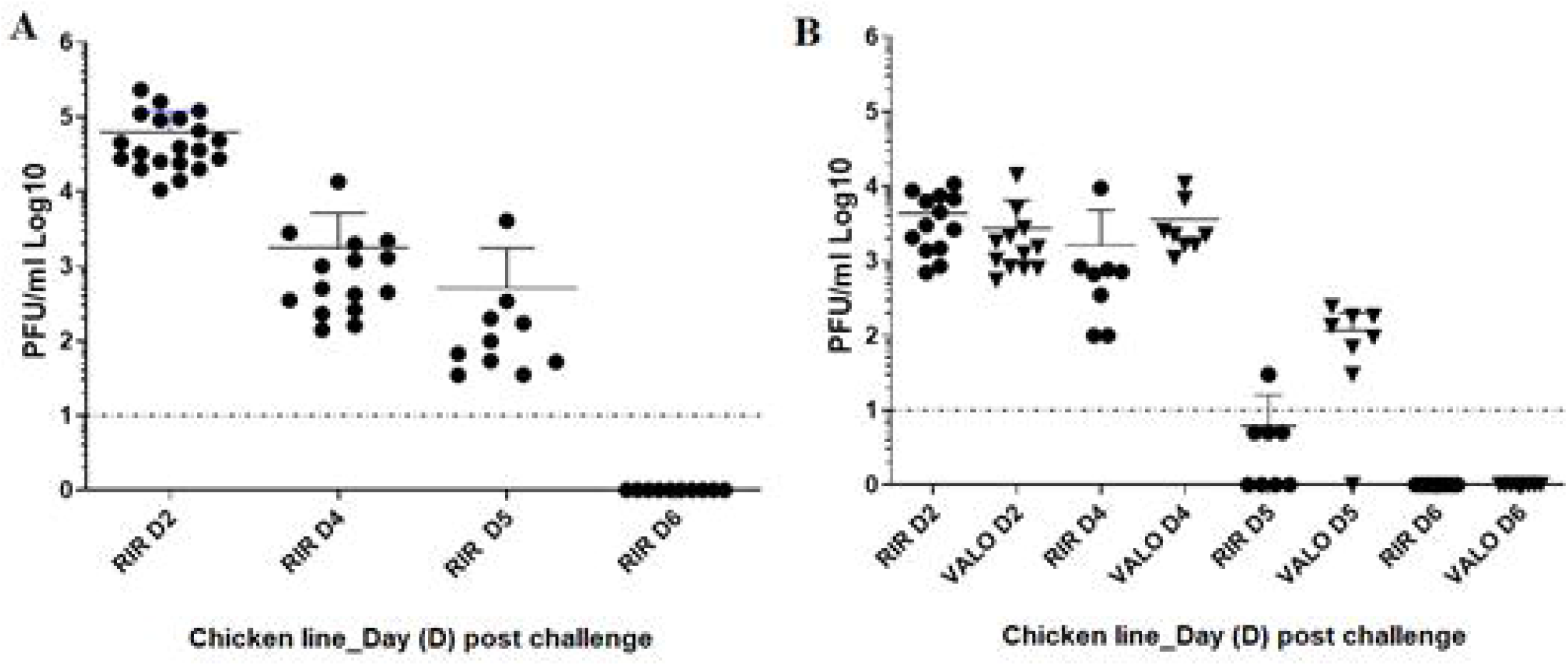
Oropharyngeal virus shedding from chickens after challenge with H9N2 AIV A/chicken/Pakistan/UDL01/08. (A). Viral titre recovered from oral swabs at day 2 (D2), 4 (D4), 5 (D5), 6 (D6) post AIV challenge from SPF RIR chickens (Experiment 1). (B). Viral titre recovered from oral swabs at day 2 (D2), 4 (D4), 5 (D5), 6 (D6) post challenge from SPF RIR and SPF VALO chickens (Experiment 2). Viral titres are expressed as log10 plaque forming units (PFU) per ml. A dashed line indicates the lower limit of detection for the plaque assay carried out on MDCK cells, 0.9 log10 PFU per ml.

Antibody responses to the H9N2 virus prior to challenge and at 10 days (Experiment 1) and 14 days (Experiment 2) post-challenge were measured by Hemagglutination inhibition (HI) assay. All infected birds in both experiments seroconverted with an average HI titre above 8 log2 (Supplementary Figure 2). No significant differences between RIR and VALO infected birds were found.

### Batch processing of colon tissue from chickens did not affect the median number of operational taxonomic units (OTUs) obtained from 16S rRNA amplicon sequencing

Following euthanasia of chickens at various time-points colon samples were aseptically sampled at *post-mortem* examination, homogenised and the microbial DNA extracted. The bacterial DNA was processed to give 16S rRNA amplicons and subjected to Illumina Miseq sequencing. We performed a pre-processing and quality check on the raw reads obtained from the Illumina Miseq platform, which removed about 20% of the sequences. All sequences were then normalized to 62,000 sequences per sample resulting in a total number of 4,094,860 OTUs with a median of 119,082.000 OTUs per sample in Experiment 1 (Batch 1), 5,970,048 of OTUs with a median of 156,783 OTUs per sample in H9N2 infected groups (Batch 2) and 8,313,621 of OTUs with a median of 173,833.5 per sample in control groups in Experiment 2 (Batch 3) (Supplementary Figure 3). This raw data was then used in our described pipelines (Material and Methods) to analyse changes in the biodiversity of the chicken colon microbiome following H9N2 infection and whether chicken breed impacted these changes.

### Microbiome Alpha diversity indices increased over time in healthy chicken colon

A temporal change in diversity measure, number of OTUs, phylogenetic diversity and Shannon diversity, were observed in control groups in both experiments (Figure 2 and Figure 4). When RIR and VALO chicken colon microbiome composition was analysed weekly (Experiment 2), we identified that statistically significant changes in alpha diversity indices in control groups occur in the time frame of 14 days, especially between day 7 and day 21 of age, suggesting that a two weeks’ period was required to see significant maturity changes in healthy colon microbiome (Supplementary Table 1). Kruskal Wallis statistical testing showed a statistically significant temporal changes in number of OTUs (the number of bacterial species detected in a sample, species richness) between day 7 and day 21, day 7 and day 28 and day 7 and 32 of age in RIR chickens and between day 7 and day 21, day 7 and day 28 of age in VALO chickens (Supplementary Table 1). Similarly, phylogenetic significant changes were seen between day 7 and day 21, day 7 and day 28 and day 7 and 32 of age and day 14 and day 28 in RIR chickens And for VALO chicken between day 7 and day 21 of age (Supplementary Table 1). Shannon index, a measure of species distribution in a sample, significantly increased between day 7 and day 21 of age in both chicken breeds suggesting a more even community evolves over time (Supplementary Table 1). Spearman correlation coefficient (rs) between time and number of detected OTUs was 0.5968 (p < 0.0001) and time and phylogenetic diversity was 0.6544 (p < 0.0001) in control RIR and VALO groups.

**Figure 2.**
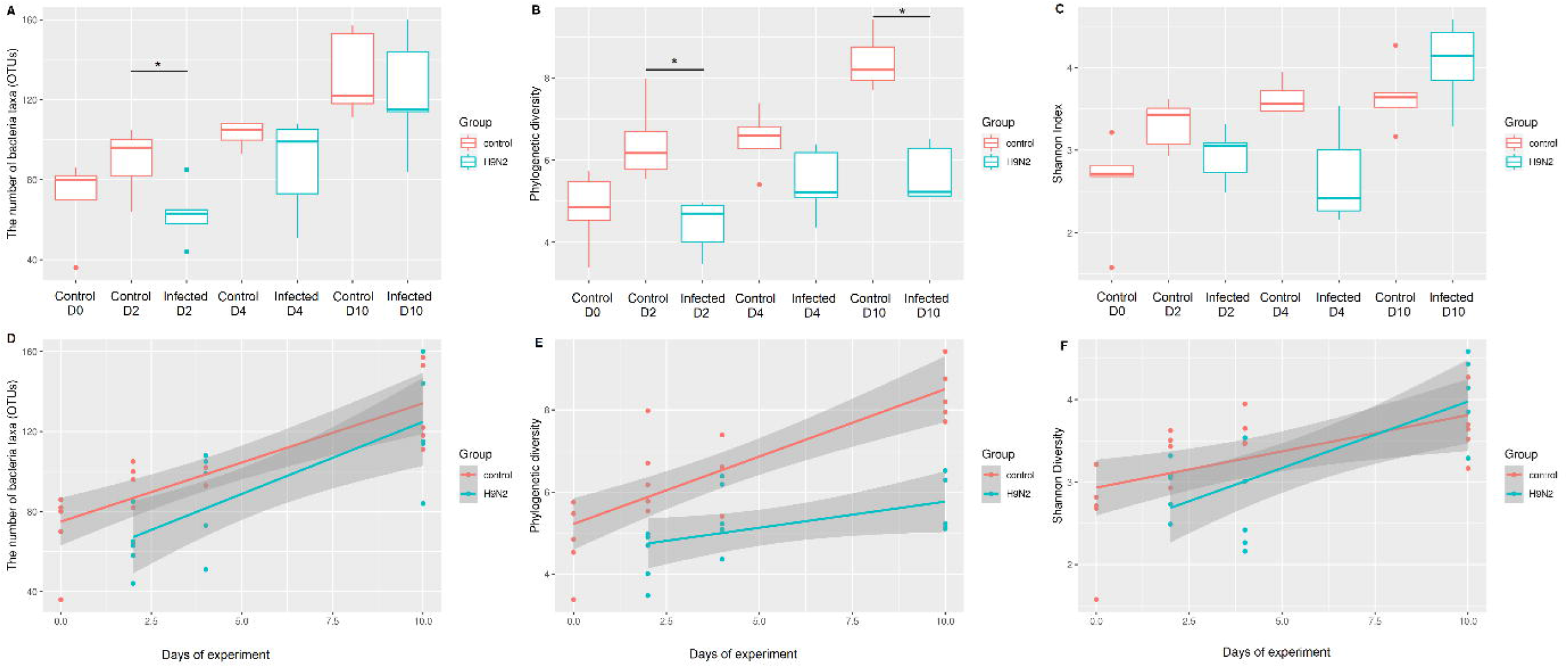
Chicken colon microbiome alpha diversity indices (A, B, C) and simple linear regression plots (D, E, F) for H9N2 infected RIR chickens compared to uninfected RIR chickens. SPF RIR chickens were challenged with H9N2 AIV A/chicken/Pakistan/UDL01/08. Colon samples were collected at day 0 (pre-challenge), day two, day four and day ten post challenge from H9N2 infected group and non-infected control groups. (A) The number of observed bacteria taxa (OTUs) in different experimental groups. (B) Faith’s phylogenetic diversity in different experimental groups. (C) Shannon indices in the different groups. Kruskal Wallis pairwise statistics were used to assess differences in community richness; * P ≤ 0.05. Only statistical differences between the groups at each time point are marked on the graph. The bottom row of linear regression plots show the change in relative abundance (D), phylogenetic diversity (E), Shannon index (F) from time t to time t+1 (y-axis) in H9N2 AIV A/chicken/Pakistan/UDL01/08 infected and control groups

The association between time and microbial diversity was also tested via simple linear regression (Supplementary Table 2), with microbial alpha diversity (the number of OTUs and phylogenetic diversity metrices) used as the dependent variable (Figure 3 and 4). In control groups, the number of OTUs, phylogenetic diversity and Shannon index significantly correlated with time (p>0.001) (Figure 2 and 4). However, individual variation within the colon microbiome, especially VALO line at later time points (day 28 and day 32 of age) was greater as compared to age of 7-or 14-days-old.

**Figure 3.**
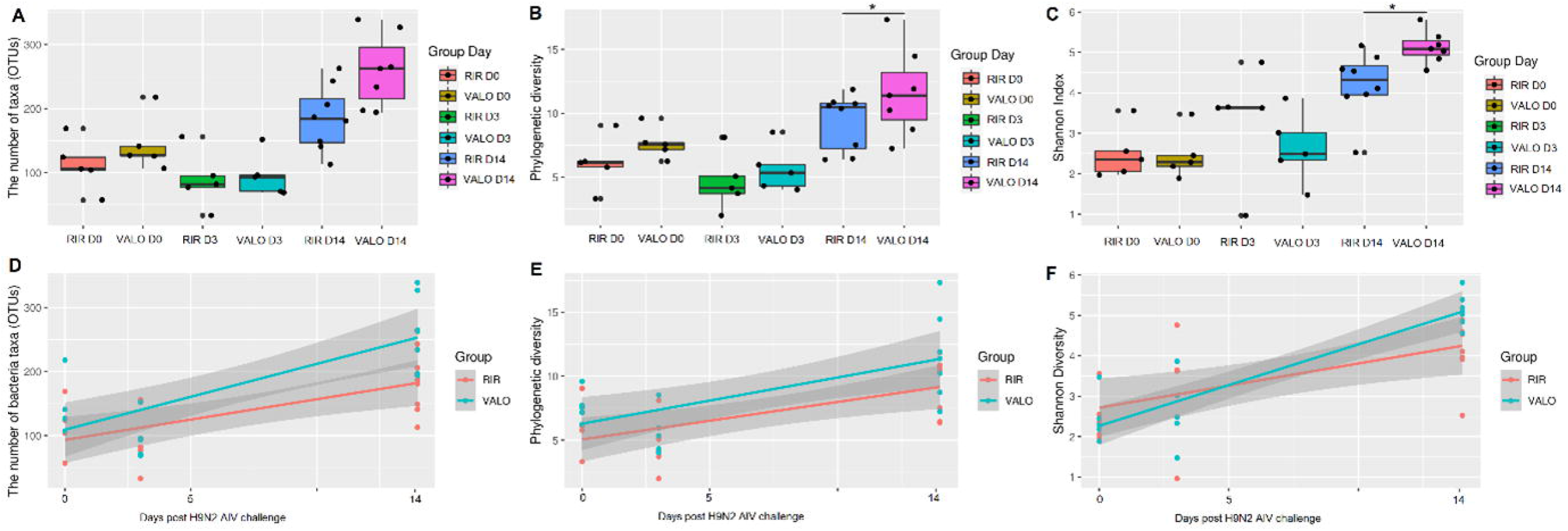
Alpha diversity indices of colon microbiome compared for two divergent chicken lines, RIR and VALO. (A) The number of observed bacteria taxa (OTUs) at different time points post infection. (B) Faith’s phylogenetic diversity at different time points post infection. (C) Shannon indices at different time points post infection. The bottom row of linear regression plots shows the change in relative abundance (D), phylogenetic diversity (E), Shannon index (F) from time t to time t+1 (y-axis). Colon samples were collected pre-challenge, at day 0 of experiment, (D0) and day 3 (D3) and day 14 post challenge (D14). Chickens were challenge with recombinant A/chicken/Pakistan/UDL01/08 H9N2 LPAIV. Kruskal Wallis pairwise statistics were used to assess differences in community richness; * P ≤ 0.05.

**Figure 4.**
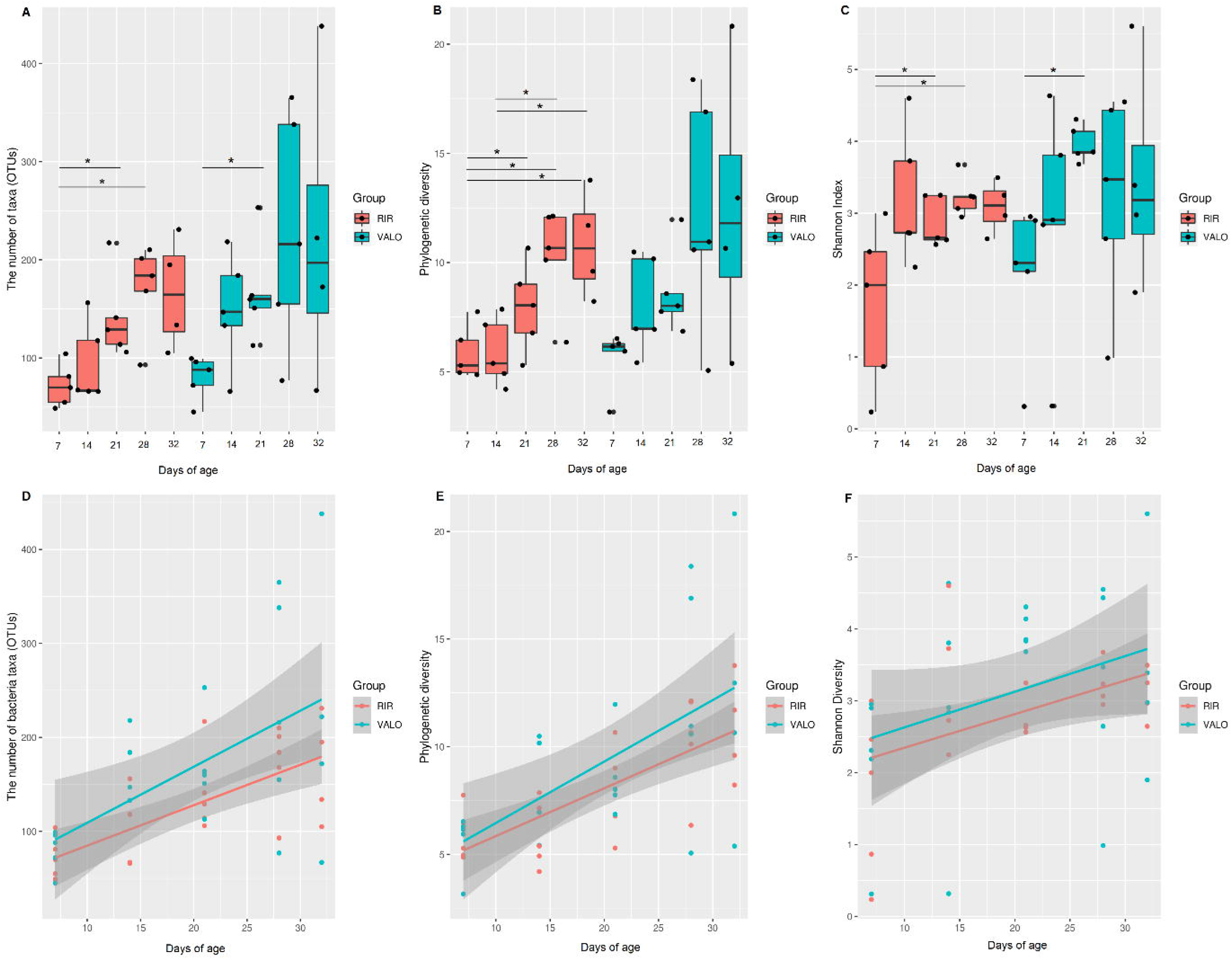
Temporal changes in RIR and VALO chicken’s colon microbiome. (A) The number of observed bacteria taxa (OTUs) at different time points. (B) Faith’s phylogenetic diversity at different time. (C) Shannon indices at different time points. Kruskal Wallis pairwise statistics were used to assess differences in community richness; * P ≤ 0.05. The bottom row of linear regression plots shows the change in relative abundance (D), phylogenetic diversity (E), Shannon index (F) from time t to time t+1 (y-axis) over the time in two divergent chicken lines. Colon samples were collected at day 7 of age, day 14, day 21, day 28 and day 32 of age. Major changes in alpha diversity indices were seen in 14 days interval, especially between day 7 and day 21 of age.

### Colon microbiome alpha diversity indices are significantly lower during the acute phase of H9N2 infection in chickens which is not altered in two different chicken breeds

First alpha diversity was quantified by the total number of observed species (OTUs) in each sample in experiment 1 (H9N2 infected versus sham infected RIR chickens) (Figure 2A). We observed a significant reduction (p<0.05 as determined Kruskal Wallis testing), in OTU number in colon samples from chickens at 2 days post influenza virus challenge (Figure 2A and Supplementary Table 3), which corresponds with high levels of viral shedding (Figure 1A), compared to the sham infected group at the same timepoint. The OTU numbers at day four and ten post challenge were not significantly different to the control group however, we observed higher variability in interquartile range of OTU numbers with the infected groups at days four and ten compared to the corresponding control groups and compared to the day 2 groups. This suggests that the acute reduction in number of OTU on day two occurred more uniformly than the recovery of OTU numbers post infection in individuals (Figure 2A). Faith’s phylogenetic index of observed species which describes OTU diversity and Shannon diversity index which indicates OTU evenness were also measured (Figure 2B and 2C). Kruskal Wallis statistical testing showed statistically significant (p < 0.01) reduced phylogenetic alpha diversity in H9N2 infected RIR chicken at acute phase of infection (day 2 post-challenge) and at day 10 post challenge as compared to control groups at the same time point measured (Figure 2B and Supplementary Table 3). We observed no statistical difference in the Shannon Index between any of the infected groups and associated controls (Figure 2C and Supplementary Table 3). Simple linear regression analysis showed that the Shannon index and the number of OTUs increased over the course of H9N2 infection in infected RIR group, similarly to the control group (Figures. 2D, 2F and Supplementary Table 2). The number of OTUs significantly correlated with time, as determined by linear regression (R2=0.666, p <0.0001, equation Y = 5.914*X + 74.92 for control RIR chickens, and R2 =0.510, p= 0.0004, equation Y = 5.914*X + 62.44 for H9N2 infected RIR chickens) (Figure 2D and Supplementary Table 2). The Faith’s phylogenetic diversity significantly correlated with time for both the control and infected group but the increase in phylogenetic diversity of the infected group was retarded as compared to the control group, suggesting that diversity development of the chicken colon microbiome was reduced by H9N2 virus infection (R^2^= 0.6894, p <0.000, equation Y = 0.3286*X + 5.220 for control group, and R^2^= 0.2114, p = 0.0414, equation Y = 0.1057*X + 4.653 for the H9N2 infected group) (Figure 2E and Supplementary Table 2).

Figure 3 shows the alpha diversity measurements compared for RIR and VALO chicken breeds infected with H9N2 at day zero, day three and day fourteen post challenge. As it was seen in experiment 1, the number of OTUs and phylogenetic diversity dropped during the acute phase of infection (day 3 post infection) following H9N2 AIV challenge in both chicken lines (Figure 3A and 3B) however this was not statistically significant (Supplementary Table 3). Between the chicken breeds no statistically significant differences in alpha diversity metrices before the challenge (day zero, D0) and during the acute phase of infection (day 3 post-challenge) (Figure 3) were observed, both lines responded in a similar fashion to infection by H9N2 AIV. Interestingly, an increased Faith’s phylogenetic (Figure 3B) and Shannon diversity (Figure 3C) indices were found in VALO chickens as compare to RIR chickens at recovery phase of H9N2 infection (day 14 post-challenge) (p < 0.05) (Supplementary Table 3).

### Beta diversity gut community changes are associated with H9N2 AIV infection and chicken breed

To compare the beta diversity among the groups at different time points, we performed Principal Coordinates Analysis (PCoA) and Principal component analysis (PCA) using the unweighted Unifrac data of taxonomic composition that includes phylogenetic diversity metrics (Figure 5). A significant separation in the control groups was observed over the time of birds’ maturity (Figure 5A). Significant differences in beta diversity within the RIR and VALO breed control groups were found in time-based manner of at least 7 days interval. Analysis of variance have shown significant differences between day 7 and 14, day 21 and 32, day 14 and 21, day 28 and 32 but not between day 28 and 32 in both chicken breed control groups (Supplementary Table 4). The only statistically significant difference between RIR and VALO chicken breeds was seen at day 21 of age for the control groups (Figure 5A and Supplementary Table 4).

**Figure 5.**
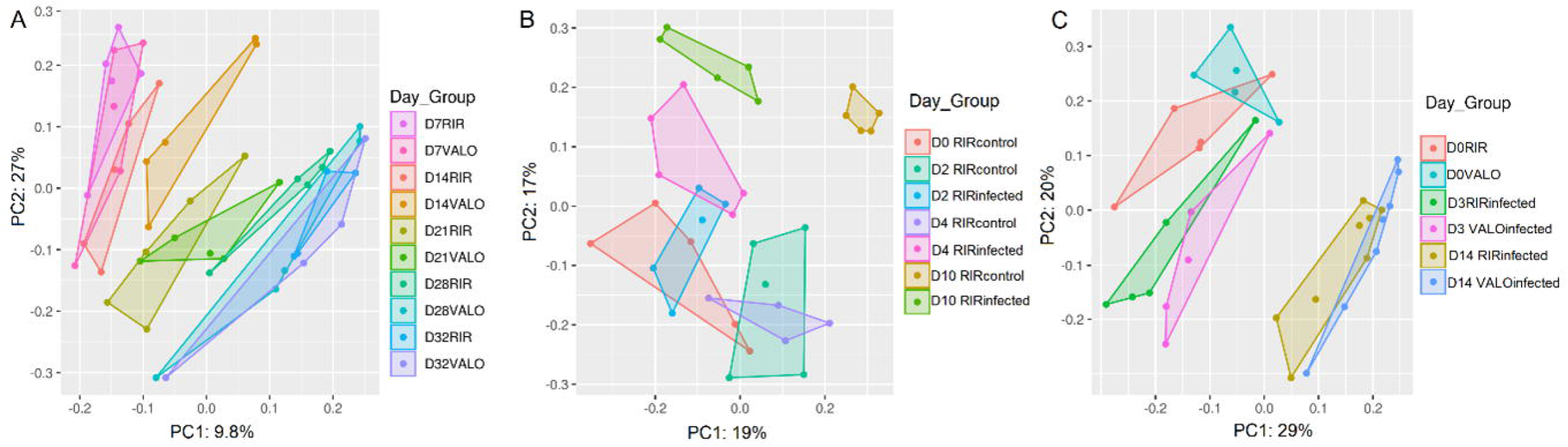
Compositional principal coordinate analysis (PCA) plot of chicken colon microbiome using unweighted UniFrac distance data, categorized according to the time points and groups (Day_Group). (A). PCA analysis of RIR and VALO chicken colon microbiome grouped by day. The samples were collected at day 7 of age (D7), day 14 (D14), day 21 (D21), day 28 (D28) and day 32 of age (D32). (B). PCA analysis of H9N2 infected and uninfected, control RIR chickens at day 0 (D0) pre-challenge, day 2 (D2), day 4 (D4) and day 10 post-challenge (D10). Chickens were challenge with recombinant A/chicken/Pakistan/UDL01/08 H9N2 LPAIV at D0 of experiment. (C). PCA analysis of H9N2 infected RIR and VALO chickens at day 0 pre-challenge (D0), day 3 (D3) and day 14 (D14) post-challenge. Chickens were challenge with recombinant A/chicken/Pakistan/UDL01/08 H9N2 LPAIV at D0 of experiment. PC1, PC2; percent variables explained (%).

PCoA plots indicate a significant separation between control and H9N2 RIR infected chickens at all time points tested (day two, day four, and day ten post challenge) for experiment 1 (Figure 5B). Analysis of variance (PERMANOVA) for measuring beta-diversity showed that the H9N2 RIR infected group had significantly lower diversity as compared to RIR control group at all time points tested (Supplementary Table 4). Similarly, significant separation in beta diversity was observed between day zero and day three post challenge for both RIR and VALO H9N2 infected chickens (Figure 5C). Analysis of variance showed significant lower diversity in RIR and VALO infected groups as compared to sham infected groups at the same time point tested (Supplementary. Table 4).

### Bacterial taxa associated with H9N2 infection

A mean relative abundance of the dominant bacteria at phyla, class, order, and family levels between H9N2 AIV RIR infected and control chickens (Experiment 1) is shown in Supplementary Figure 4 whereas between RIR and VALO H9N2 infected chickens and its corresponding controls (Experiment 2) is shown in Supplementary Figure 5. Analysis of composition of microbiomes (ANCOM) was applied against group and day of infection variables to determine which bacteria were significantly differentiated in relative abundance at genus level (Figure 6). The ANCOM results showed significant differences between the control and H9N2 infected RIR groups in members of the *Furmicutes* phylum. Six of *Furmicutes* phylum, *Peptostreptococcaceae* (*Terrisporobacter*), *Planococcaceae* (*Lysinibacillus*), *Erysipelotrichaceae* (*Turicibacter*), *Lachnospiraceae* (*Cellulosilyticum*), *Paenibacillacea* (*Paenibacillus*), *Clostridiaceae* 1 (*Clostridium sensu stricto* 1), were significantly different between the RIR control and H9N2 infected groups and had a high W-statistics and f-score (Figure 6A). Detailed significant statistics of ANCOM percentile of different taxa is shown in (Supplementary Table 5). Furthermore, we also performed Linear discriminant analysis Effect Size (Lefse) analysis based on OTUs to compare the microbial communities between RIR control and RIR H9N2 infected birds at each time point tested. The LEfse analysis and ANCOM generated similar results (Figure 6 and Figure 7). LEfse results indicated differences in the phylogenetic distributions of the microbiome of H9N2 infected and control chickens at the OTU level (Figure 7). The gut microbial communities in H9N2 infected birds were different compared to those in control groups. A histogram of the LDA scores was computed for features that showed differential abundance between H9N2 infected and control chickens (Figure 7A and 7B). The LDA scores indicated that the relative abundances of Streptococcaecae (*Streptococcus*), and Planococcaceae (*Lysinibacillus*) were much more enriched in H9N2 infected birds versus control at day 2 post-challenge (Figure 7A) and the most differentially abundant bacteria taxa (LDA score [log 10] > 3). The most differentially abundant bacterial taxon in control birds was characterized by a preponderance of Peptostreptococcaceae (LDA score [log10] > 3) at day 2 post-challenge (Figure 7A). The differences in the phylogenetic distributions of the microbiomes of H9N2 infected and control chickens at the OTU level were also found at day 4 post-challenge (Figure 7B). The LDA scores indicated that the relative abundances of Penicibacillacea (Penicibacillus), Planococcaceae (Lysinibacillus), Erysipelotrichaceae (Turicibacter), Clostridiaceae (*Clostridium sensu stricto* 1) were much more enriched in H9N2 infected birds as compared to control birds at day 4 post-challenge (Figure 7B). The control birds were characterized mainly by a preponderance of different Clostridia and Bacillaceae (Bacillus) (LDA score [log10] > 3). In addition, we saw differential abundance of bacterial taxa between H9N2 infected RIR and VALO and their relative control groups in our second experiment (Figure 8). The LDA scores indicated that the relative abundances of Clostridiales (*Clostridium sensu stricto* 1) and Planococcaceae (Lysinibacillus) (LDA score [log10] > 3.5) were much more enriched in RIR infected birds at day 3 post challenge as compared to the control group, and this correspond to our results obtained in first experiment at day 4 post-challenge (Figure 7B). In VALO infected chicken, LDA scores indicated that the relative abundances of Aerococcaceae, Paenibacillaceae, Bacillaceae (LDA score [log10] > 3.5) were much more enriched at day 3 post-challenge whereas Clostridiales, Peptostreptococcae and Brevibacteriacea (LDA score [log10] > 3.5) were significantly reduced in the VALO infected group compared to control group at day 3 post challenge (Figure 8).

**Figure 6.**
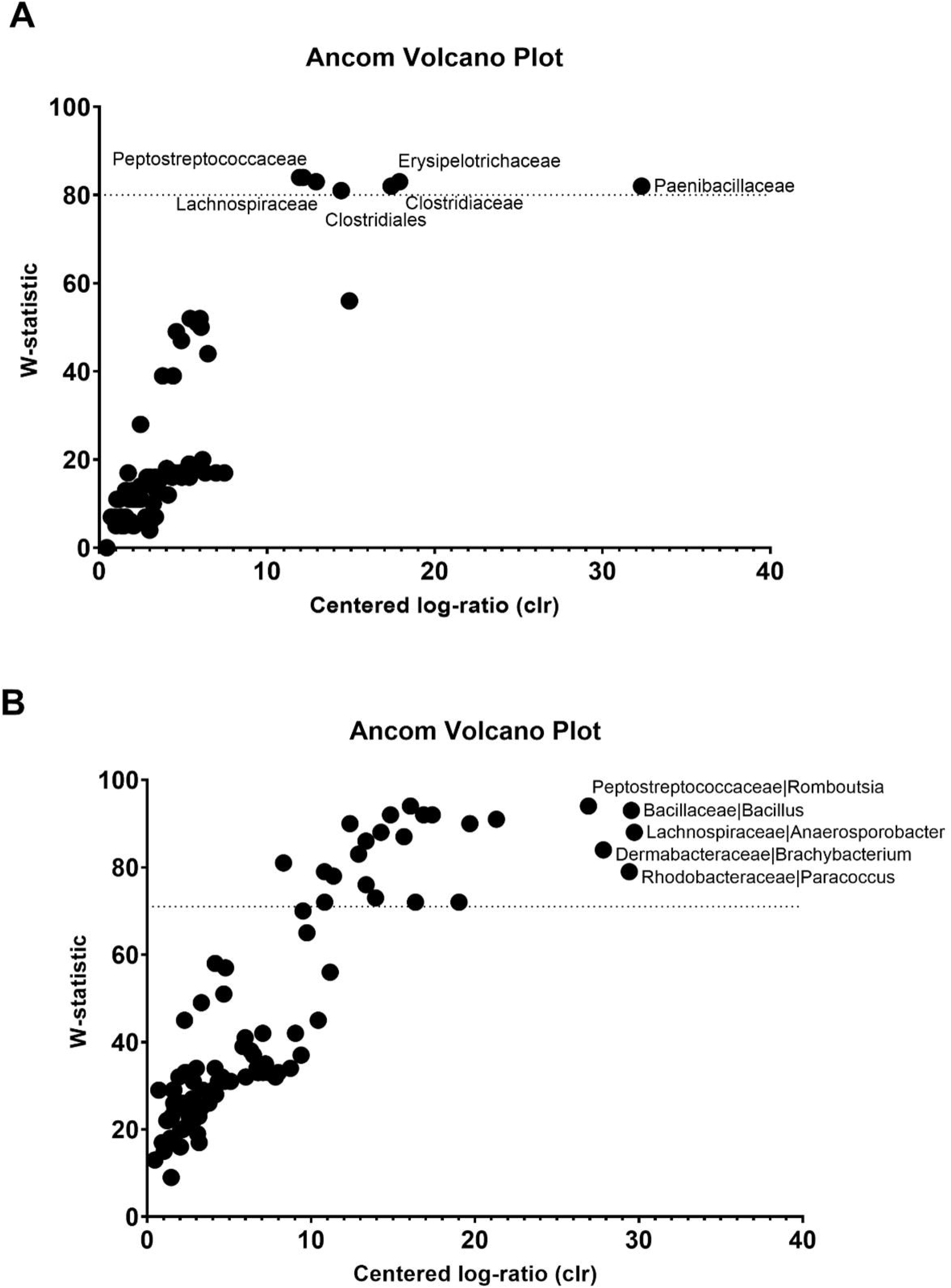
Volcano plot for the analysis of composition of microbiomes (ANCOM) test. Significant bacterial taxa are above the line. Taxa on the top-left corner are distinct species but with small proportions, (low f-score). Truly different taxa are depicted as one moves towards the far right (high W-statistic). A. ANCOME test applied to control RIR and H9N2 infected RIR chickens (Experiment 1). B. ANCOME test applied to RIR and VALO H9N2 infected chickens (Experiment 2).

**Figure 7.**
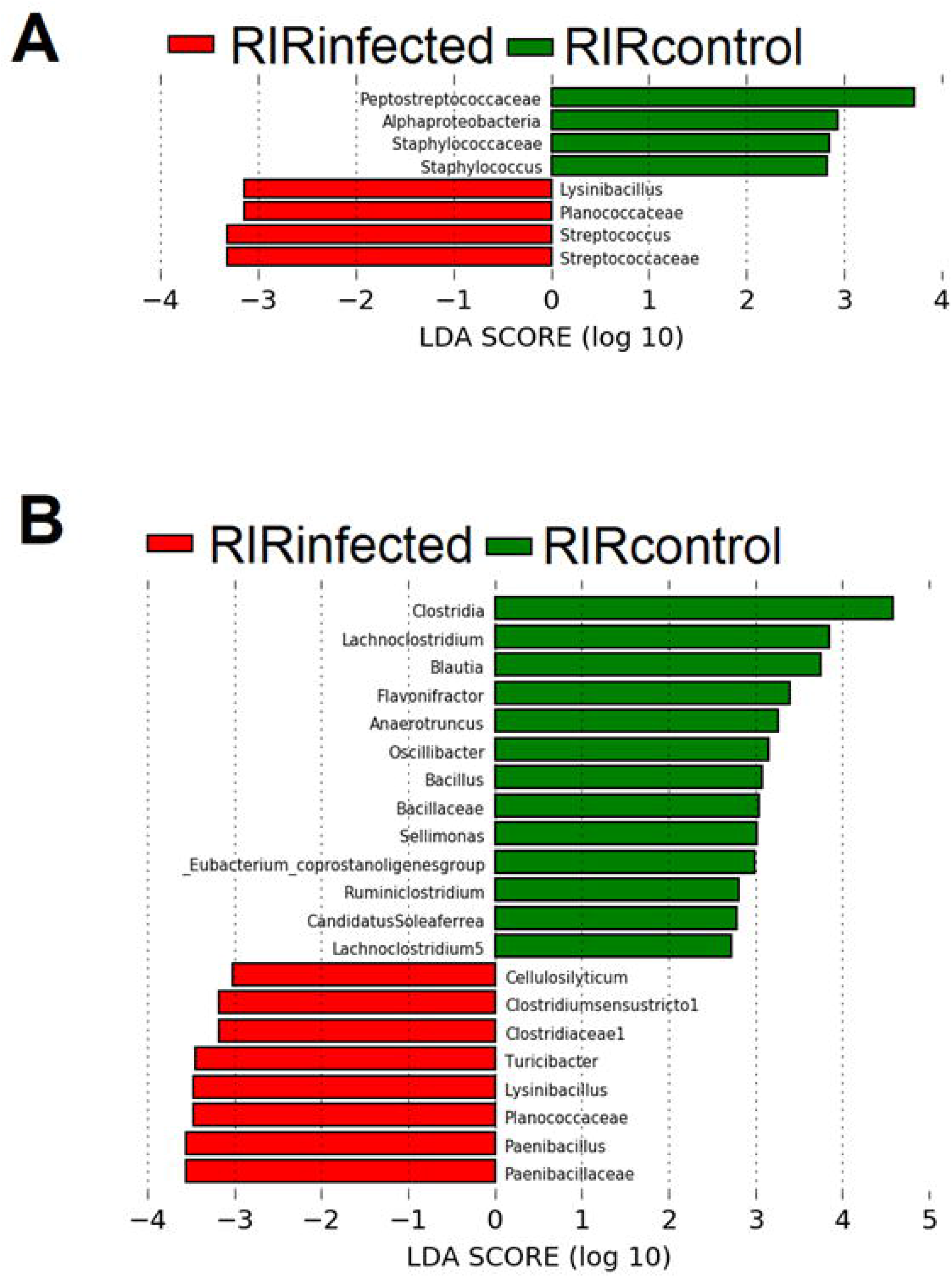
Linear discriminant analysis Effect Size (LEfSe) analysis identifying taxonomic differences in the colon microbiota of RIR H9N2 AIV infected and control chickens. (**A**) Cladogram using the LEfSe method indicating the phylogenetic distribution of colon microbiota associated with H9N2 infection in RIR chickens and control group at day 2 post-challenge. Histogram of LDA scores of 16S gene sequences in H9N2 infected chickens at day 2 (**A**) and at day 4 post challenge (**B**) and respective control groups. LDA scores (log_10_) above 3.0 and *P* <0.05 are shown. Chickens were challenge with A/chicken/Pakistan/UDL01/08 H9N2 LPAIV.

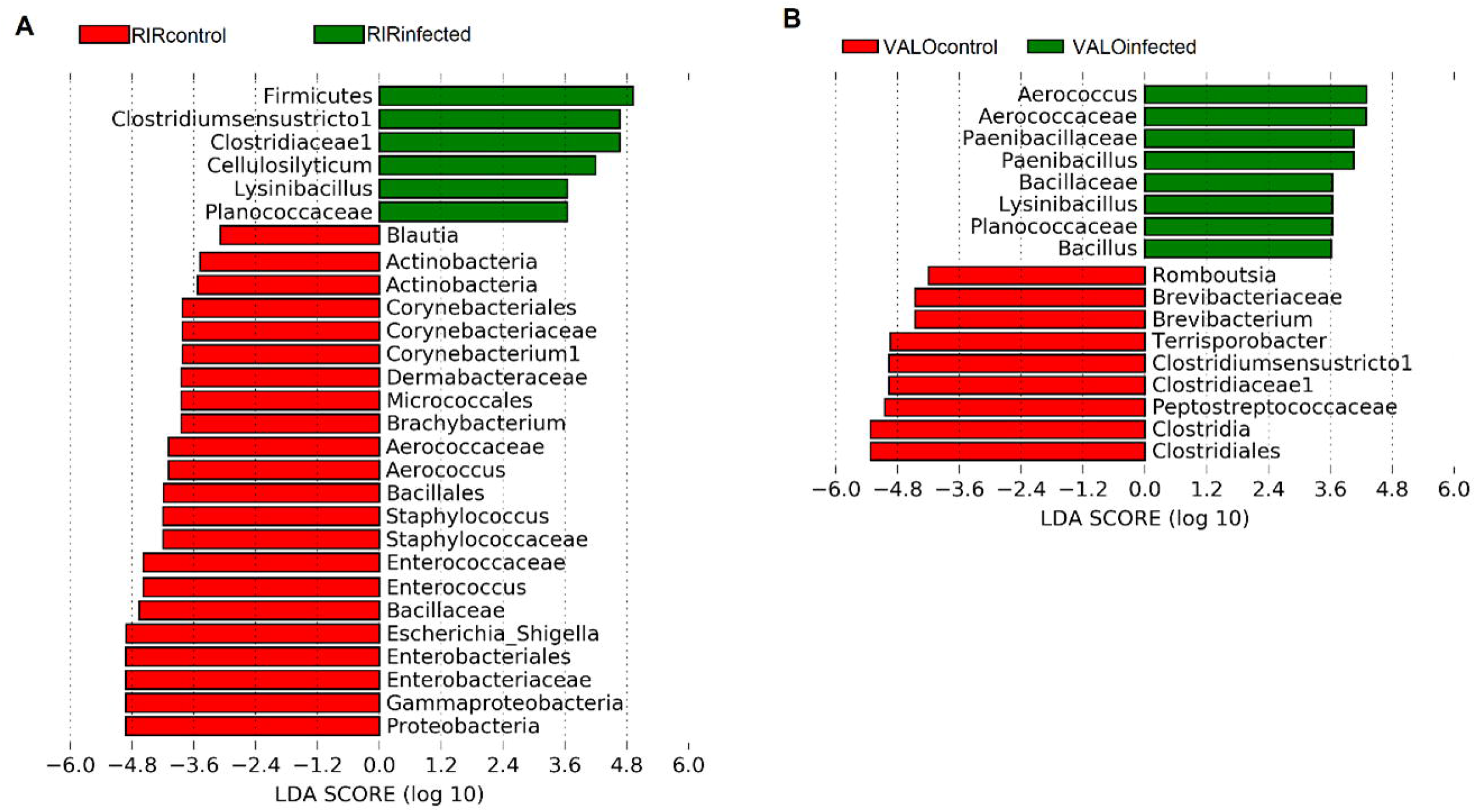

## Discussion

A tremendous number of microorganisms (bacteria, viruses, and fungi), collectively termed the microbiome, are associated with the various host mucosal surfaces and play an important role in host homeostasis [28, 43, 44]. Those microorganisms undergo dynamic changes due to numerous factors, including ageing, changes in diet, environment, or infection by pathogens [28, 45–47]. Recent studies have shown that interaction between host microbiome and viruses may play a crucial role in dictating disease pathogenesis in mammalian hosts [24, 48, 49]. As in all vertebrates, chicken mucosal surfaces are share by diverse and dynamic population of microbiome [50–52]. In this study, we observed that chicken gut microbiome diversity changes during host maturity, from day 7 post hatch to day 32 post hatch (Figure 4). The alpha diversity, measured as the number of bacterial taxa, phylogenetic diversity and Shannon index in healthy colon increased over the time of chicken growth in both chicken breeds (RIR and VALO), with most changes being seen in 14 days interval. Temporal increases in the Shannon index suggests a more even bacterial microbiome community evolving with age. Like our findings, Xi, et al. [53] showed that major changes in chicken gut microbiome development were observed between day 14 and day 28 post hatch. We found that beta diversity also strongly correlates with time, with the major changes observed in 7 days intervals. Although ageing shaped both alpha and beta diversity in chickens, beta diversity changes occur more rapidly than alpha. The only statistically significant difference in beta diversity of the gut microbiome between RIR and VALO chicken breeds was seen at day 21 of age (Figure 4).

H9N2 AIV infection in chickens occurs via the respiratory route and the predominant site of initial viral replication is mucosal surface of the oropharyngeal cavity, followed by infection and replication in other sites of the respiratory and intestinal tracts [13, 54]. H9N2 infection in chickens clinically manifests with nonspecific symptoms [55], we observed a mild lethargy and diarrhoea especially between day 2 and day 4 post-challenge in infected chickens of both breeds. Neither breed lost body weight or failed to gain body weight in comparison to the uninfected control groups following H9N2 AIV challenge, Supplementary Figure 1, suggesting that infected chicken consume similar amount of feed as their controls and that this behaviour does not account for the changes in microbiome composition following challenge as it has observed in the influenza infected mice model [56]. Both breeds shed the virus via the oropharyngeal route, but not via cloaca which agrees with other LPAIV infection studies where little or no shedding was observed from cloacal cavity (Figure 1) [57–59].

Although we showed in this study that the number of bacterial taxa and Faith’s phylogenetic diversity index significantly correlated with time, this correlation was impaired by H9N2 AIV infection. H9N2 infection decreased the number of bacteria taxa during the acute phase of infection (peak viral shedding) and phylogenetic diversity at both acute and recovery phase of infection (Figures 2 & 3). This suggests that the colon microbiome of H9N2 AIV infected birds lost it overall richness at the acute phase of infection and the microbiome reconstruction appears via increased numbers of predominant bacteria resulting in greater microbiological evenness but not via increased phylogenetic diversity compared to control birds. Zhao, et al. (2018) have shown that species richness of faecal microbiome in swans positive for H5N1 AIV infection tended to be lower than that in healthy controls [60]. Furthermore, it was shown that approximately 1,100 of the OTUs identified in the healthy-control swan samples were not detected in AIV H5N1-positive samples [60]. Hird, et al. (2018) have shown that overall species richness in duck species infected with AIV differs within the duck species and for instance only *Anas platyrhynchos* and *Anas carolinensis* showed a significant decrease in alpha diversity in the AIV positive individual [61]. In the current study, we also showed increased Shannon index and phylogenetic diversity in VALO infected chicken as compared to RIR at day 14 post challenge that might suggest different microbiome recovery dynamics occurred between two chicken breeds (Figure 3). Furthermore, we also observed that beta diversity changes are associated with H9N2 AIV infection at all time points measured in this study (Figure 5). Like our study, differences in microbiome composition between the control group and H9N2 infected chickens was seen in ileal contents [30], cecum content [29] and faecal swab samples [62] by others. In contrast, Zhao, et al. (2018) did not notice beta diversity changes in faecal swab samples obtained from migrating whooper swans infected with H5N1 influenza virus [60]. Furthermore, Hird, et al. [2018] have shown that the microbiome may have a unique relationship with influenza virus infection at the species level [61]. All those findings might suggest that relationship between host microbiome and influenza virus infection might depends on host genetic background of the avian species that is infected by AIV [61] and the strain of infecting AIV. However, these associations require further evaluation.

In this study, taxonomic analysis showed significant changes in diversity and abundance the healthy colon chicken followed H9N2 LPAIV infection regardless the chicken line infected (Figures 7 & 8). Healthy chicken colon in both chicken lines was characterized by predominance of *Proteobacteria* phylum (Enterobacteriales order) and *Firmicutes* phylum (Lactobacillales, Bacillales and Clostridiales orders) and the major differences between H9N2 infected and non-infected chickens were seen in *Firmicutes* phylum. In general, RIR chicken microbiome at acute phase of H9N2 LPAIV, expanded Bacillales, among the others. Specifically, Bacillales (*Lysinibacillus*) and Lactobacilales (Streptocococeae) (at day 2 post infection), Clostidiales (*Clostridium sensu stricto* 1) and Bacillales (*Lysinibacilus*) at day 3 post infection and Bacillales (*Lysinibacillus* and *Penbacillus*) at day 4 post infection. The question why Bacillales (*Lysinibacillus*) or Streptocococeae are overrepresented during acute phase of influenza infection is open and needs further evaluation.

In the current study, we observed a dynamic change in chicken colon between acute and recovery phase of AIV infection suggesting that different bacteria taxa might play a role in recovery from infection. Interestingly, phylogenetic abundance distributions of the microbiome in chicken colon differ substantially at day 3 post challenge between RIR and VALO infected chickens (Clostridiales in RIR versus Bacillales and Enterobacteriales in VALO) but was similar at recovery (day 14 post challenge) in both chicken lines, mostly represented by high abundance of Bacillales and Clostridiales (*Clostriudium sensu stricto* 1). At the same time healthy colon was enriched by Clostridiales (Peptostreptococcae) in RIR and Lactobacillales in both chicken lines. It was previously shown in mice model that *Lactobacillus* spp. have probiotic potential and can improve immune control in influenza infected individuals and thus could aim microbiome recovery following infection [63–65].

In conclusion, we have shown for the first-time dysbiosis in healthy colon microbiome following AIV of H9N2 subtype in two divergent chicken lines. We observed significantly reduced alpha and beta biodiversity during infection. Although bird breed did not influence the general trend observed in alpha diversity, it has an impact on beta diversity. *Firmicutes* phylum was the most differentially abundant between infected and non-infected individuals. Lactobacillales was missing at recovery phase of infection for both breeds suggesting supplementation of this taxa in during recovery could be beneficial. Predominance of different bacteria taxa at different time points during influenza infection suggests there is an involvement of chicken gut microbiome. Modulating the composition of the gut microbiome using probiotics might serve to promote a healthy community that confers protection or mitigates disease from influenza virus infection in chickens.

## Materials and Methods

### Ethics Statement

All animal work was approved and regulated by the UK government Home Office under the project license (P68D44CF4) and reviewed by the Pirbright Animal Welfare and Ethics Review Board (AWERB). All personnel involved in the procedures were licensed by the UK Home Office.

### In vivo chicken study design

Two separate *in vivo* experiments were performed. In Experiment 1, we used specific pathogen free (SPF) Rhode Island Red (RIR) chickens to assess the effect on H9N2 infection on RIR host colon microbiome. A total of 45 one-day-old SPF RIR chickens were host in pens until 2-weeks of age when chickens were randomly allocated into two experimental groups: control (n=25) and H9N2 challenged (n=20). Five control group birds were culled before the challenge at 3-weeks of age to establish the starting microbiome profile. Colon samples were collected at day 2 (n=10), day 4 (n=10) and day 10 (n=20) post challenge from both infected and control groups. In Experiment 2, SPF RIR were compared to SPF VALO chickens to assess differences in host microbiome following H9N2 infection in the two different chicken breeds. A total of 104 one-day-old SPF chickens (n=52, RIR and n=52, VALO) were host separately in two pens. Colon samples were collected from each breed control group (n=5, RIR and n=5, VALO) at day 7, 14, 21, 28 and 32 of age. At three weeks of age, 18 birds from each group were randomly selected and H9N2 challenged. The colon samples from challenged birds were collected before the challenge (day 0), at day 3 post-challenge and day 14 post-challenge.

RIR chickens were provided as day old chicks from the National Avian Resource Facility (NARF) located at The Roslin Institute, Edinburgh, UK whilst VALO chickens were delivered as fertilised eggs from VALO BioMedia GmbH (Germany) which were set and hatched at the Biological Service Unit (Poultry) at The Pirbright Institute (UK). The feed was provided *ad libitum* according to manufacture instruction for the chicken age. Both RIR and VALO chicks move from starter feed to grower at 3 weeks old. In both experiments AIV challenged chickens were housed in self-contained BioFlex^®^ B50 Rigid Body Poultry isolators (Bell Isolation Systems) maintained at negative pressure. The H9N2 challenged birds received 100μl of 10^4^ pfu (50μl in each nare) of H9N2 AIV, A/chicken/Pakistan/UDL01/08. Blood samples were taken from a wing vein pre-challenge and at day 14 post-challenge for serum collection. All birds were swabbed daily from day of challenge until 8 days post infection in both cloacal and buccal cavities to determine viral shedding. Swabbing was carried out with sterile polyester tipped swabs (Fisher Scientific, UK) which were transferred into viral transport media (Who, 2006), vortexed briefly, clarified by centrifugation and stored at −80 °C prior to virus detection. At appropriate timepoints chickens were humanly euthanized either by intravenous administration of sodium pentobarbital if housed in isolators or by cervical dislocation in the case of the control groups. All colon samples were collected from the distal part of colon (2-cm sections of each chicken), and then snap-frozen. Samples were stored at −80°C until subsequent analysis. Body weights were monitored daily until the end of experiment.

### Virus and cells

Recombinant A/chicken/Pakistan/UDL01/08 H9N2 virus was generated using reverse genetics as previously described [66]. Virus stocks were produced via passage in 10 day old embryonated chicken eggs; the allantoic fluid harvested after 48 hours and titrated by plaque assay on MDCK cells (ATCC).

Madin-Darby Canine Kidney (MDCK) cells (ATCC) were maintained in DMEM (Gibco-Invitrogen, Inc.) supplemented with 10% foetal bovine serum (Biosera, Inc.), 1% penicillin/streptomycin (Sigma-Aldrich, Inc.) and 1% non-essential aa (Sigma-Aldrich, Inc.).

### Serology

Haemagglutinin inhibition (HI) assays were carried out using challenge virus A/Chicken/Pakistan/UDL01/08(H9N2) antigen. HI assays were performed according to standard procedures [68]. Titres were expressed as log2 geometric mean titres (GMT). Samples with titres below 3 log2 GMT were considered negative.

### Virus shedding

Buccal swab samples from both challenge experiments were titrated by plaque assay on MDCK cells. MDCKs were inoculated with 10-fold serially diluted samples and overlaid with 0.6% agarose (Oxoid) in supplemented DMEM (1× MEM, 0.21% BSA V, 1 mM L-Glutamate, 0.15% Sodium Bicarbonate, 10 mM Hepes, 1× Penicillin/Streptomycin (all Gibco) and 0.01% Dextran DEAE (Sigma-Aldrich, Inc.), with 2 μg/ml TPCK trypsin (SIGMA). They were then incubated at 37 °C for 72 h. Plaques were developed using crystal violet stain containing methanol. Viral titres were expressed as log10 plaque forming units (PFU) per ml and the limit of detection is 0.9 log10 PFU per ml for this assay.

Cloacal swab samples from both challenge experiments were titrated by qRT-PCR assay for the viral matrix (M) protein. qRT-PCR analysis was completed using the Superscript III Platinum one-step qRT-PCR kit (Life Technologies). Cycling conditions were: (i) 5-min at 50°C, (ii) a 2-mins step at 95°C, and (iii) 40 cycles of 3 sec at 95°C, 30 s of annealing and extension at 60°C. Cycle threshold (CT) values were obtained using 7500 software v2.3. Mean CT values were calculated from triplicate data. Within viral M segment qRT-PCR, an M segment RNA standard curve was completed alongside the samples to quantify the amount of M gene RNA within the sample from the CT value. T7 RNA polymerase-derived transcripts from UDL-01 segment 7 were used for preparation of the standard curve.

### DNA extraction and 16S rRNA gene amplification

Samples were extracted in batches (experiment 1, one batch and experiment 2, two batches). DNA was extracted using the PowerSoil^®^ DNA Isolation Kit (Mo Bio) according to manufacturer instruction. DNA extraction reagent only controls were included for each batch of DNA extractions along with ZymoBIOMICS Microbial Community Standards (Zymo Research) and E. coli DH5α (ThermoFisher). The V2-V3 region of the 16S rRNA gene was amplified via PCR as described previously by Glendinning et al. (2016) [51]. Briefly, a nested PCR protocol was performed using the V1-V4 primers 28F (‘5–175 GAGTTTGATCNTGGCTCAG-3’) and 805R (‘5-GACTACCAGGGTATCTAATC-3’) followed by the V2-V3 primers 104F (‘5-GGCGVACGGGTGAGTAA-3’) and 519R (‘5–177 GTNTTACNGCGGCKGCTG-3’) with Illumina adaptor sequences and barcodes.

### Sequencing and data analysis

Libraries were analysed on a High Sensitivity DNA Chip on the Bioanalyzer (Agilent Technologies) and Qubit dsDNA HS assay (Invitrogen). The amplicon libraries were pooled in equimolar concentrations, before loading on the flow cell of the 500 cycle MiSeq Reagent Kit v2 (Illumina, USA) and pair-end sequencing (2 × 250 bp). The amplicon-based sequencing was performed using the Illumina MiSeq platform at The Pirbright Institute. Bioinformatic analysis was implemented using the Quantitative Insights into Microbial Ecology (QIIME) platform version qiime2-2019.10. Low-quality sequencing reads were quality trimmed and denoise using DADA2. Potential chimeric sequences were removed using UCHIME, and the remaining reads assigned to 16S rRNA operational taxonomic units (OTUs) based on 97% nucleotide similarity with the UCLUST algorithm and then classified taxonomically using the SILVA reference database (silva-132-99-nb-classifier). Taxonomy was then collapsed to the genus-level. The microbial community structure was estimated by microbial biodiversity (i.e., species richness and between-sample diversity). Shannon index, phylogenetic diversity, and the obseof age whilst RIR and VALO AIVrved number of species were used to evaluate alpha diversity, and the unweighted UniFrac distances were used to evaluate beta diversity. All these indices (alpha and beta diversity) were calculated by the QIIME pipeline. Data was visualized using R package “ggplot2” ver 3.2.1 [67].

### Statistical analysis

Kruskal Wallis pairwise statistics were used to assess differences in community richness (Shannon diversity, phylogenetic diversity, and the observed number of species, OTU). In addition, Spearman correlation coefficient and simple linear regression was used to evaluate temporal changes in community richness that occurred during AIV infection between control and infected birds. A multivariate ANOVA (PERMANOVA) analysis was used to determine significant differences in β diversity distances across groups. Principal-coordinate analysis (PCoA) graphs were constructed to visualize similarity between the samples over the time of AIV infection. Additionally, Principal Component Analysis (PCA) was performed using OTU matrix. The Linear Discriminant Analysis Effect Size (LEfSe) algorithm and analysis of composition of microbiomes (ANCOME) were used to identify differentially abundant taxa between the groups. For LEfSe analysis, depends on the experiments, different groups were assigned as comparison classes and were analysed by days. Briefly, in experiment 1, RIR control and RIR AIV infected groups were assigned as comparison classes and assessed at day 0, day 2, day 4 and day 10 post-challenge. In experiment 2, RIR control and VALO control groups were assigned as comparison classes and analysed at 7, 14, 21, 28 and 32 day of age whilst RIR and VALO AIV infected groups that represented separate classes were analysed at day 0, day 3 and day 14 post-challenge. LEfSe identified features that were statistically different between assigned groups and then compared the features using the non-parametric factorial Kruskal-Wallis sum-rank test (alpha value of 0.05) and Linear Discriminant Analysis (LDA) >2.0.

## Funding Information

This work described herein was funded by The Pirbright Institute BBSRC ISP grant BBS/E/I/00007030, BBS/E/I/00007034, BBS/E/I/00007038 and BBS/E/I/00007039.

## Acknowledgments

We would like to acknowledge colleagues at The Pirbright Institute who supported the *in vivo* work; Elizabeth Billington, Dr Jean-Remy Sadeyen, Professor Munir Iqbal and the poultry unit team and Illumina MiSeq sequencing; Dr Graham Freimanis.

## Author contributions

The work was conceptualized by HS and KC. Experimental work was executed by KC, MZ, DB, JL, RLR, KC, and HS. The Manuscript was written by KC and HS and edited by all authors.

## Notes

### Competing Interest Statement

The authors have declared no competing interest.

