## supplementary figures for "Low pathogenic avian influenza virus infection retards colon microbiome diversification in two different chicken lines"

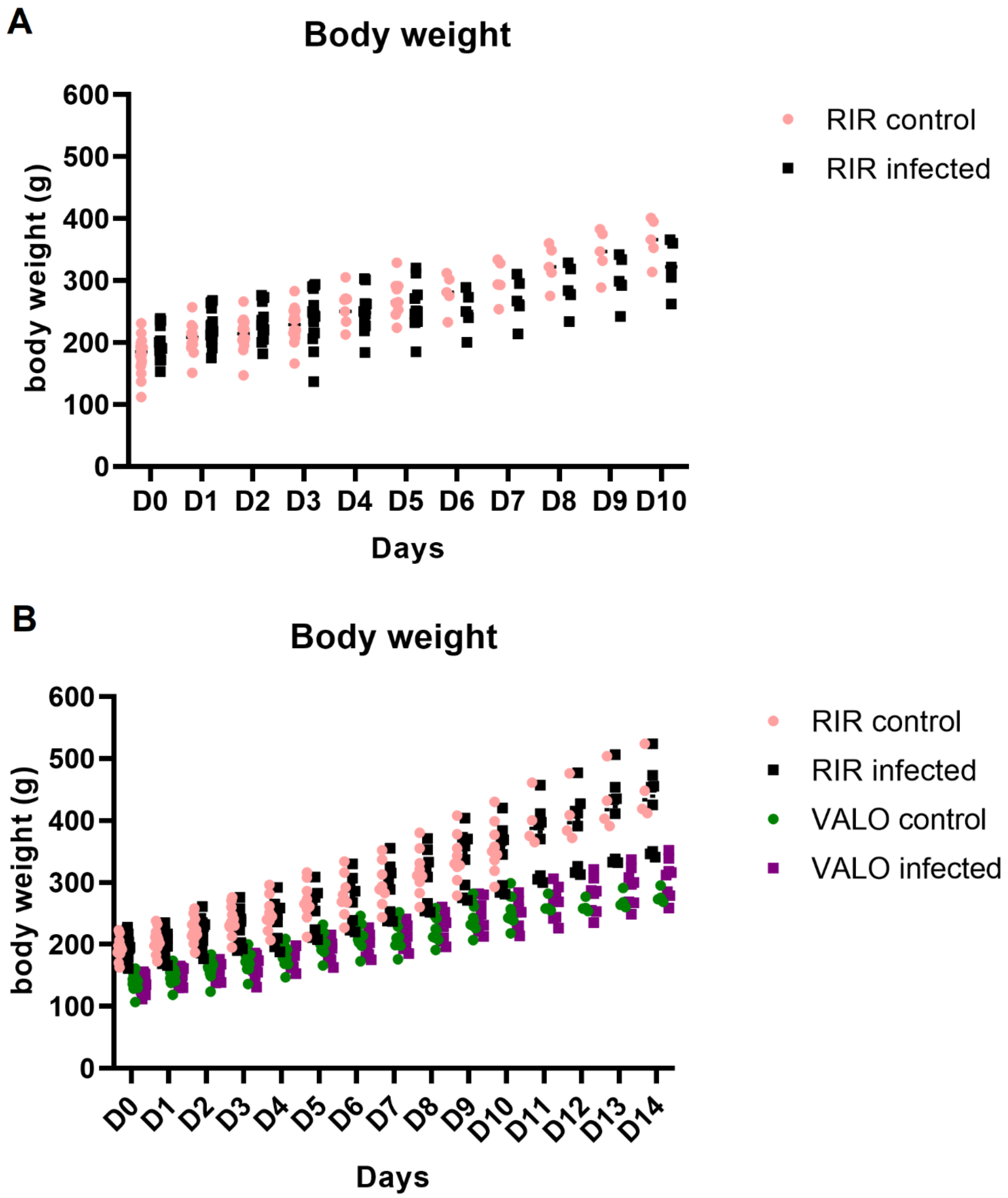


**Suppl. Fig. 1. Body weight (g) of the chickens during influenza virus infection period.** Two separate experiments were performed where all chickens were weighed daily from day of infection (D0) until 14 days post challenge (D14). (A) In experiment 1, SPF Rhode Island Red

(RIR) chickens were either challenged with LPAIV A/chicken/Pakistan/UDL01/08(H9N2) (RIR infected) or remained unchallenged (RIR control). (B) In experiment 2, SPF RIR chickens (RIR control and infected) and SPF VALO chickens (VALO control and infected) were challenge with LPAIV A/chicken/Pakistan/UDL01/08(H9N2).

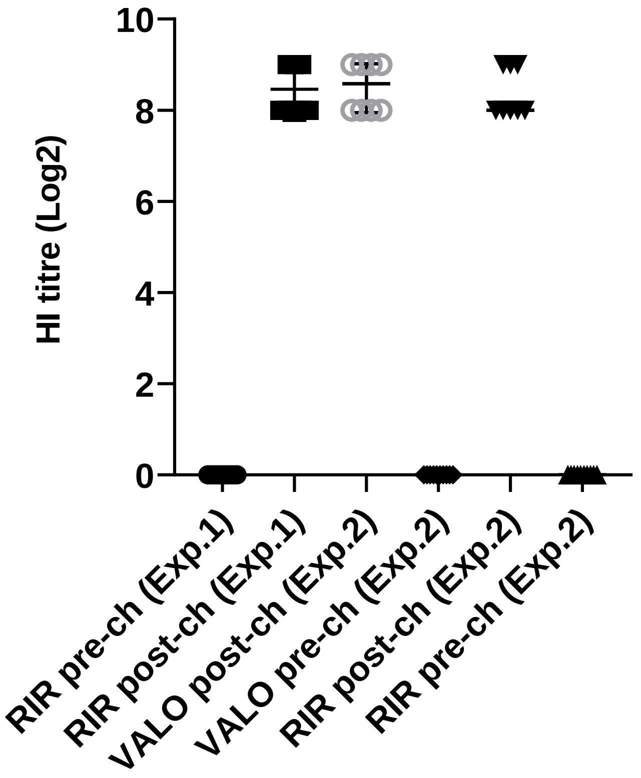


**Suppl. Figure 2. HI titers pre- and post-challenge with the LPAIV A/chicken/Pakistan/UDL01/08(H9N2).** For the two separate experiments (Exp. 1 and Exp.2) blood collected was collected from each AIV challenged bird prior to infection (pre-ch) and at post challenge (post-ch). Sera was harvested from the individual blood samples and used to perform HI assays to assay for H9N2 antibodies. Individual HI titers (log2) are plotted and standard error for groups of at day 0 (pre-ch) and day 10 (Experiment 1) or 14 (Experiment 2) post-challenge.

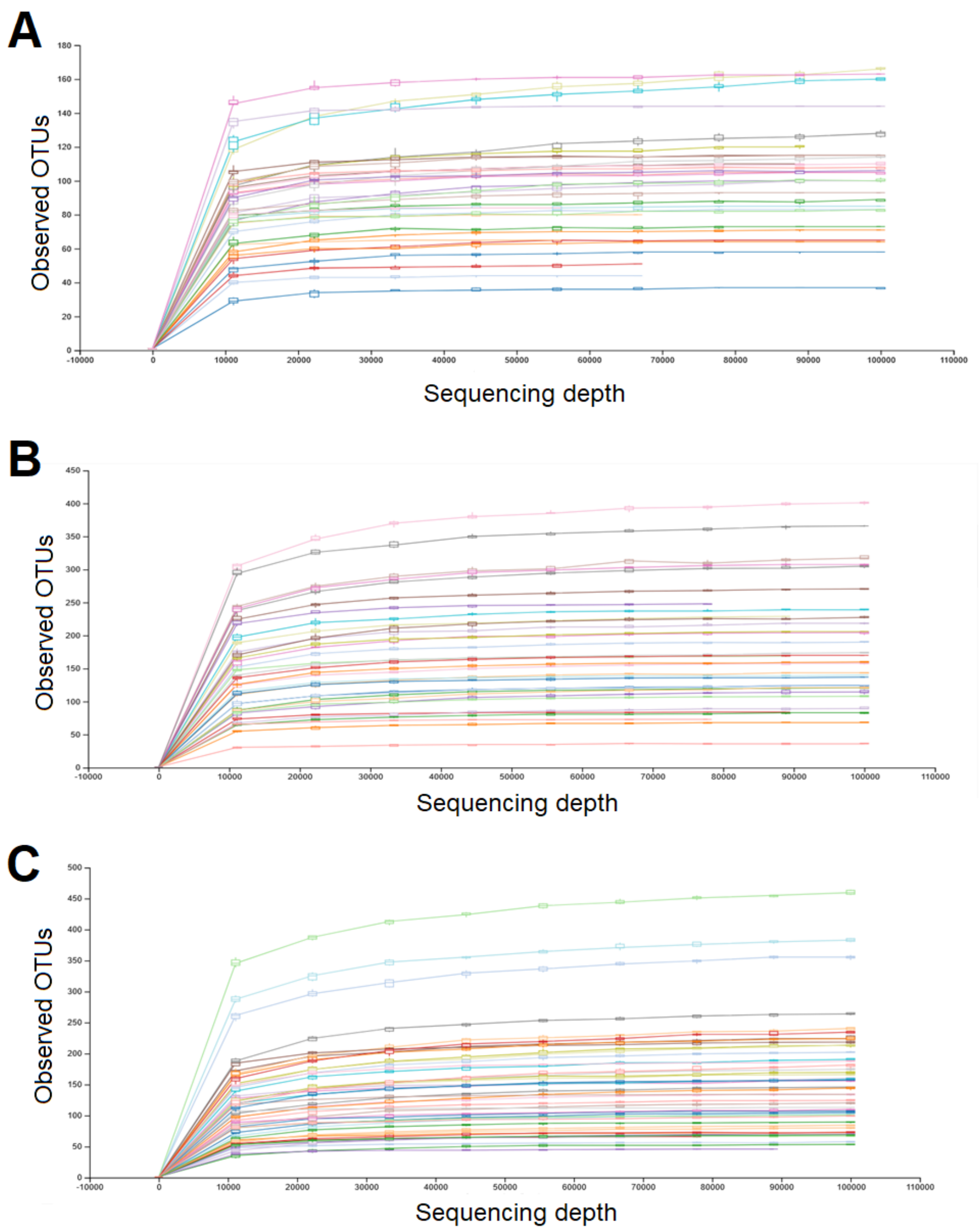


**Suppl. Figure 3**. Rarefaction analysis for the observed species. Rarefaction curve for each individual microbiome samples obtained from: (A) RIR and RIR H9N2 infected chickens (B) RIR and VALO H9N2 infected chickens. (C) RIR and VALO control chickens. Curves were plotted at a cut-off of 0.03 for each sample. All samples were normalized to 62,000 sequences per sample for further analysis.

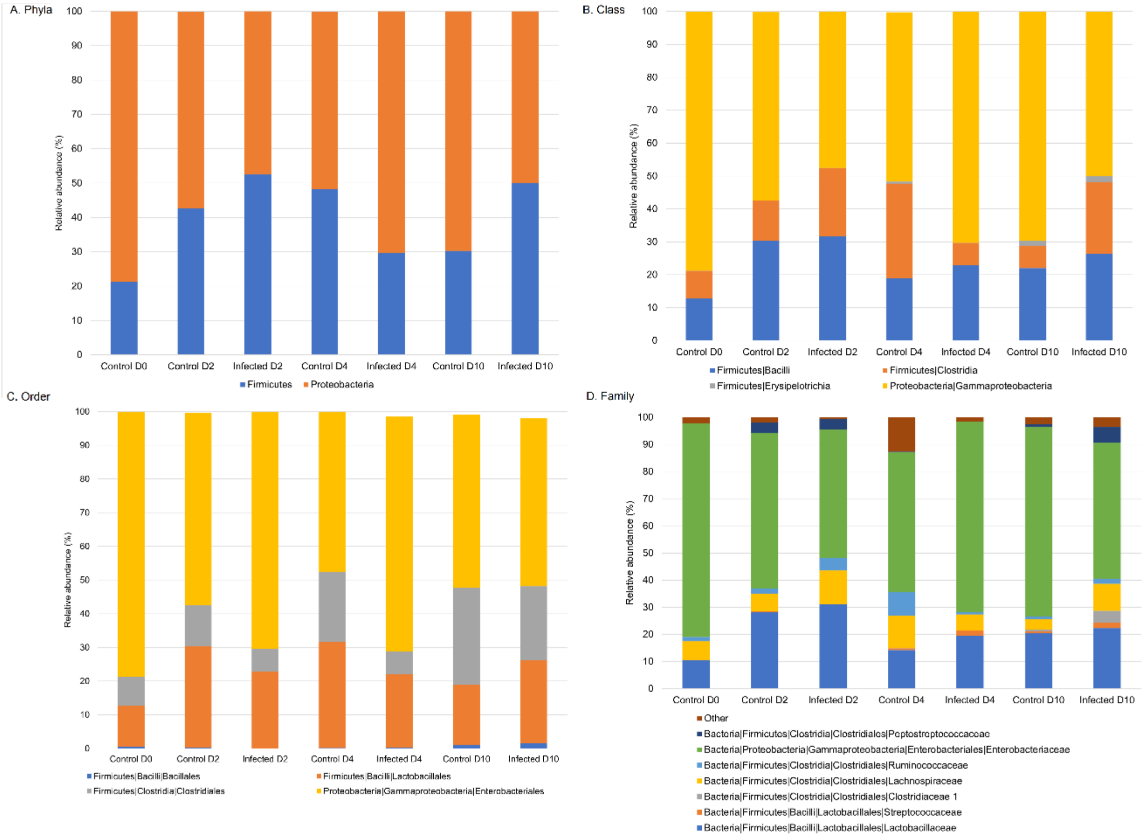


**Suppl. Figure 4**. Mean relative abundance of the dominant bacterial phyla (A), class (B), order (C), and family (D) of SPF RIR control and SPF RIR H9N2 AIV infected chickens at different time points testes. All the bacteria taxa which accounted for less than 1% of the total sequences are placed in artificial category "Other". The colon samples were collected at day 0 (D0) pre-challenge, day 2 (D2), day 4 (D4) and day 10 post-challenge (D10). Chickens were challenge with recombinant A/chicken/Pakistan/UDL01/08 H9N2 LPAIV at D0 of experiment.

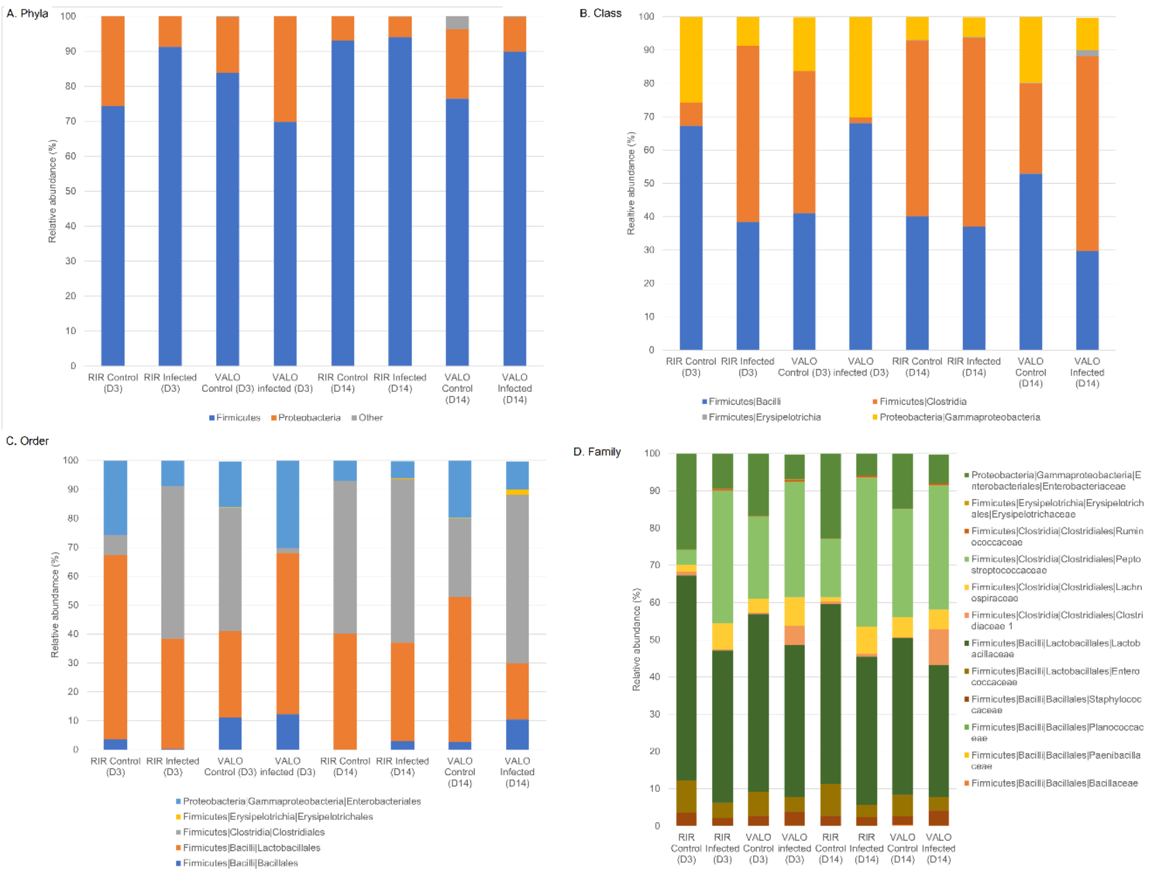


**Suppl. Figure 5**. Mean relative abundance of the dominant bacterial phyla (A), class (B), order (C), and family (D) of SPF RIR and VALO H9N2 AIV infected chickens SPF RIR and VALO and its corresponding controls at different time points testes. All the bacteria taxa which accounted for less than 1% of the total sequences are placed in artificial category “Other”. The colon samples were collected at day 0 (D0) pre-challenge, day 3 (D3) and day 14 post-challenge (D14). Chickens were challenge with recombinant A/chicken/Pakistan/UDL01/08 H9N2 LPAIV at D0 of experiment.
