## supplementary figures for "Low pathogenic avian influenza virus infection retards colon microbiome diversification in two different chicken lines"

Supplementary Tables

Supplementary Table 1. Kruskal-Wallis H tests of associations between time and colon microbiota composition in healthy chickens. Kruskal-Wallis test on alpha diversity indices was conducted to examine the differences in microbiota composition that occurred over the time in two divergent chicken lines. In Experiment 1, specific pathogen free (SPF) Rhode Island Red (RIR) chickens were used to assess changes in host colon microbiota. Samples were taken at day 0 (D0), day 4 (D4), day 10 (D10) of experiment. In Experiment 2, SPF RIR were compared to SPF VALO chickens. Samples were collected in 7-days interval, at day 7 of age (D7), day 14 of age (D14), etc. OTUs, the number of bacteria taxa.

| Alpha diversity | Group 1 | Group 2 | H | p-value | q-value |
|---|---|---|---|---|---|
| | | **Experiment 1** | | | |
| OTUs | | | | | |
| | RIR D0 | RIR D2 | 2.15122 | 0.142457 | 0.186974 |
| | | RIR D4 | 6.05042 | 0.013903 | 0.038018 |
| | | RIR D10 | 6.818182 | 0.009023 | 0.038018 |
| | RIR D2 | RIR D4 | 2.178151 | 0.139983 | 0.186974 |
| | | RIR D10 | 6.818182 | 0.009023 | 0.038018 |
| | RIR D4 | RIR D10 | 6.05042 | 0.013903 | 0.038018 |
| Faith`s Index | | | | | |
| | RIR D0 | RIR D2 | 1.843636 | 0.174525 | 0.21559 |
| | | RIR D4 | 3.84 | 0.050044 | 0.075065 |
| | | RIR D10 | 3.938182 | 0.047202 | 0.075065 |
| | RIR D2 | RIR D4 | 3.84 | 0.050044 | 0.075065 |
| | | RIR D10 | 4.810909 | 0.02828 | 0.059388 |
| | RIR D4 | RIR D10 | 6 | 0.014306 | 0.037553 |
| Shannon Index | | | | | |
| | RIR D0 | RIR D2 | 4.810909 | 0.02828 | 0.065987 |
| | | RIR D4 | 6 | 0.014306 | 0.048881 |
| | | RIR D10 | 5.770909 | 0.016294 | 0.048881 |
| | RIR D2 | RIR D4 | 2.16 | 0.141645 | 0.198303 |
| | | RIR D10 | 3.152727 | 0.0758 | 0.13265 |
| | RIR D4 | RIR D10 | 0.06 | 0.806496 | 0.846821 |
| | | **Experiment 2** | | | |
| OTUs | | | | | |
| | D7 RIR | D14 RIR | 3.152727 | 0.0758 | 0.284251 |
| | | D21 RIR | 3.938182 | 0.047202 | 0.225196 |
| | | D28 RIR | 5.770909 | 0.016294 | 0.104745 |
| | | D32 RIR | 3.84 | 0.050044 | 0.225196 |
| | D14 RIR | D21 RIR | 0.534545 | 0.464702 | 0.746843 |

| | | | | | |
|---|---|---|---|---|---|
| | | D28 RIR | 0.272727 | 0.601508 | 0.802551 |
| | | D32 RIR | 0 | 1 | 1 |
| | D21 RIR | D28 RIR | 0.883636 | 0.347208 | 0.624974 |
| | | D32 RIR | 1.5 | 0.220671 | 0.472867 |
| | D28 RIR | D32 RIR | 0.06 | 0.806496 | 0.930572 |
| | D7 VALO | D14 VALO | 1.32 | 0.250592 | 0.490289 |
| | | D21 VALO | 6.818182 | 0.009023 | 0.101514 |
| | | D28 VALO | 1.843636 | 0.174525 | 0.413349 |
| | | D32 VALO | 2.16 | 0.141645 | 0.413349 |
| | D14 VALO | D21 VALO | 1.843636 | 0.174525 | 0.413349 |
| | | D28 VALO | 0.010909 | 0.916815 | 1 |
| | | D32 VALO | 0.24 | 0.624206 | 0.802551 |
| | D21 VALO | D28 VALO | 0.272727 | 0.601508 | 0.802551 |
| | | D32 VALO | 1.5 | 0.220671 | 0.472867 |
| | D28 VALO | D32 VALO | 0 | 1 | 1 |
| Faith`s Index | | | | | |
| | D7 RIR | D14 RIR | 0.010909 | 0.916815 | 0.937652 |
| | | D21 RIR | 3.938182 | 0.047202 | 0.173228 |
| | | D28 RIR | 4.810909 | 0.02828 | 0.159076 |
| | | D32 RIR | 6 | 0.014306 | 0.122202 |
| | D14 RIR | D21 RIR | 2.454545 | 0.117185 | 0.239697 |
| | | D28 RIR | 4.810909 | 0.02828 | 0.159076 |
| | | D32 RIR | 6 | 0.014306 | 0.122202 |
| | D21 RIR | D28 RIR | 2.454545 | 0.117185 | 0.239697 |
| | | D32 RIR | 2.94 | 0.086411 | 0.204657 |
| | D28 RIR | D32 RIR | 0 | 1 | 1 |
| | D7 VALO | D14 VALO | 3.152727 | 0.0758 | 0.200648 |
| | | D21 VALO | 6.818182 | 0.009023 | 0.122202 |
| | | D28 VALO | 3.152727 | 0.0758 | 0.200648 |
| | | D32 VALO | 2.16 | 0.141645 | 0.25496 |
| | D14 VALO | D21 VALO | 0.272727 | 0.601508 | 0.75917 |
| | | D28 VALO | 2.454545 | 0.117185 | 0.239697 |
| | | D32 VALO | 1.5 | 0.220671 | 0.331007 |
| | D21 VALO | D28 VALO | 1.32 | 0.250592 | 0.352395 |
| | D21 VALO | D32 VALO | 0.96 | 0.327187 | 0.446164 |
| | D28 VALO | D32 VALO | 0.06 | 0.806496 | 0.907308 |
| Shannon Index | | | | | |
| | D7 RIR | D14 RIR | 3.152727 | 0.0758 | 0.284251 |
| | | D21 RIR | 3.938182 | 0.047202 | 0.225196 |
| | | D28 RIR | 5.770909 | 0.016294 | 0.104745 |
| | | D32 RIR | 3.84 | 0.050044 | 0.225196 |
| | D14 RIR | D21 RIR | 0.534545 | 0.464702 | 0.746843 |
| | | D28 RIR | 0.272727 | 0.601508 | 0.802551 |
| | | D32 RIR | 0 | 1 | 1 |
| | D21 RIR | D28 RIR | 0.883636 | 0.347208 | 0.624974 |
| | D21 RIR | D32 RIR | 1.5 | 0.220671 | 0.472867 |

| | | | | |
|---|---|---|---|---|
| D28 RIR | D32 RIR | 0.06 | 0.806496 | 0.930572 |
| D7 VALO | D14 VALO | 1.32 | 0.250592 | 0.490289 |
| | D21 VALO | 6.818182 | 0.009023 | 0.101514 |
| | D28 VALO | 1.843636 | 0.174525 | 0.413349 |
| | D32 VALO | 2.16 | 0.141645 | 0.413349 |
| D14 VALO | D21 VALO | 1.843636 | 0.174525 | 0.413349 |
| | D28 VALO | 0.010909 | 0.916815 | 1 |
| | D32 VALO | 0.24 | 0.624206 | 0.802551 |
| D21 VALO | D28 VALO | 0.272727 | 0.601508 | 0.802551 |
| | D32 VALO | 1.5 | 0.220671 | 0.472867 |
| D28 VALO | D32 VALO | 0 | 1 | 1 |

Supplementary Table 2. The association between time and colon microbial diversity tested via simple linear regression across the experimental groups. The alpha diversity (the number of OTUs and Faith`s index, phylogenetic diversity) metrices were used as the dependent variable. In Experiment 1, specific pathogen free (SPF) Rhode Island Red (RIR) chickens were used to assess the effect on H9N2 infection on RIR host colon microbiota. Samples were taken at day 0, pre-challenge (D0), day 4 post-challenge (D4), day 10 post-challenge (D10) of experiment in RIR control and RIR infected groups. In Experiment 2, SPF RIR were compared to SPF VALO chickens to assess differences in host microbiome following H9N2 infection in the two different chicken breeds. Samples were collected at day 0 (D0), pre-challenge, day 3 post-challenge (D3) and day 14 post-challenge(D14) in infected groups. In addition, control only RIR and VALO groups, samples were collected in 7-days interval, from at day 7 of age to day 32 of age.

| **Alpha diversity** | **Group** | **R2** | **Slope (Y value)** | **Equation** |
|---|---|---|---|---|
| The number of OTUs | | | | |
| Experiment 1 | RIR Control | 0.666 | 5.914 | Y = 5.914*X + 74.92 |
| | RIR Infected | 0.510 | 5.914 | Y = 5.914*X + 62.44 |
| Experiment 2 | RIR control | 0.4919 | 4.302 | Y = 4.302*X + 41.82 |
| | RIR infected | 0.4362 | 6.348 | Y = 6.348*X + 93.33 |
| | VALO control | 0.297 | 5.964 | Y = 5.964*X + 49.51 |
| | VALO infected | 0.596 | 10.27 | Y = 10.27*X + 109.3 |
| Phylogentic diversity | | | | |
| Experiment 1 | RIR Control | 0.6894 | 0.32 | Y = 0.3286*X + 5.220 |
| | RIR Infected | 0.2114 | 0.105 | Y = 0.1057*X + 4.653 |

| | | | | |
|---|---|---|---|---|
| Experiment 2 | RIR control | 0.5438 | 0.2223 | Y = 0.2223*X + 3.624 |
| | RIR infected | 0.4203 | 0.2934 | Y = 0.2934*X + 5.056 |
| | VALO control | 0.3508 | 0.2856 | Y = 0.2856*X + 3.598 |
| | VALO infected | 0.4326 | 0.3612 | Y = 0.3612*X + 6.291 |

Supplementary Table 3. Kruskal-Wallis H tests of associations between avian influenza virus (AIV) infection and colon microbiota composition in two divergent chicken lines. In Experiment 1, specific pathogen free (SPF) Rhode Island Red (RIR) chickens were used to assess the effect on H9N2 infection on RIR host colon microbiota. Samples were taken at day 0, pre-challenge (D0), day 4 post-challenge (D4), day 10 post-challenge (D10) of experiment In Experiment 2, SPF RIR were compared to SPF VALO chickens to assess differences in host microbiome following H9N2 infection in the two different chicken breeds. Samples were collected at day 0 (D0), pre-challenge, day 3 post-challenge (D3) and day 14 post-challenge(D14). In both *iv vivo* experiments chickens were challenge with recombinant A/chicken/Pakistan/UDL01/08 H9N2 LPAIV.

| Alpha diversity | Group 1 | Group 2 | H | p-value | q-value |
|---|---|---|---|---|---|
| | | Experiment 1 | | | |
| OTUs* | RIR Control (D0) | RIR Infected (D2) | 0.8836 | 0.3472 | 0.3838 |
| | RIR Control (D2) | RIR Infected (D2) | 3.9382 | 0.0472 | 0.0826 |
| | RIR Control (D4) | RIR Infected (D4) | 0.9931 | 0.319 | 0.3721 |
| | RIR Control (D10) | RIR Infected (D10) | 0.2727 | 0.6015 | 0.6316 |
| Faith`s Index | RIR Control (D0) | RIR Infected (D2) | 0.2727 | 0.6015 | 0.6316 |
| | RIR Control (D2) | RIR Infected (D2) | 6.8182 | 0.009 | 0.0316 |
| | RIR Control (D4) | RIR Infected (D4) | 3.84 | 0.05 | 0.0751 |
| | RIR Control (D10) | RIR Infected (D10) | 6.8182 | 0.009 | 0.0316 |
| Shannon Index | RIR Control (D0) | RIR Infected (D2) | 0.8836 | 0.3472 | 0.3838 |
| | RIR Control (D2) | RIR Infected (D2) | 2.4545 | 0.1172 | 0.1758 |
| | RIR Control (D4) | RIR Infected (D4) | 3.84 | 0.05 | 0.0955 |
| | RIR Control (D10) | RIR Infected (D10) | 1.8436 | 0.1745 | 0.2291 |
| | | Experiment 2. RIR and VALO infected chickens versus theirs corresponding controls | | | |
| OTUs | RIR Control (D3) | RIR Infected (D3) | 2.4545 | 0.1172 | 0.2051 |

| | | | | | |
|---|---|---|---|---|---|
| | VALO Control (D3) | VALO Infected (D3) | 3.1527 | 0.0758 | 0.1769 |
| | RIR Control (D14) | RIR Infected (D14) | 1.75 | 0.1859 | 0.2739 |
| | VALO Control (D14) | VALO Infected (D14) | 0.8929 | 0.3447 | 0.4387 |
| Faith`s Index | RIR Control (D3) | RIR Infected (D3) | 3.1527 | 0.0758 | 0.1728 |
| | VALO Control (D3) | VALO Infected (D3) | 3.1527 | 0.0758 | 0.1728 |
| | RIR Control (D14) | RIR Infected (D14) | 0.0357 | 0.8501 | 0.8816 |
| | VALO Control (D14) | VALO Infected (D14) | 0.1429 | 0.7055 | 0.823 |
| Shannon Index | RIR Control (D3) | RIR Infected (D3) | 2.4545 | 0.1172 | 0.215 |
| | VALO Control (D3) | VALO Infected (D3) | 3.9382 | 0.0472 | 0.1202 |
| | RIR Control (D14) | RIR Infected (D14) | 3.5714 | 0.0588 | 0.1372 |
| | VALO Control (D14) | VALO Infected (D14) | 2.2857 | 0.1306 | 0.2151 |
| Experiment 2. RIR and VALO infected chickens | | | | | |
| OTUs | RIR Control (D0) | VALO Control (D0) | 2.4545 | 0.1172 | 0.1547 |
| | RIR Control (D0) | RIR Infected (D3) | 1.32 | 0.2506 | 0.2891 |
| | VALO Control (D0) | VALO Infected (D3) | 3.1527 | 0.0758 | 0.1137 |
| | RIR Infected (D3) | VALO Infected (D3) | 0.0109 | 0.9168 | 0.9168 |
| | RIR Infected (D3) | RIR Infected (D14) | 6.1929 | 0.0128 | 0.0321 |
| | VALO Infected (D3) | VALO Infected (D14) | 8.0769 | 0.0045 | 0.0224 |

| | | | | | |
|---|---|---|---|---|---|
| | RIR Infected (D14) | VALO Infected (D14) | 5.1017 | 0.0239 | 0.0448 |
| Faith`s Index | RIR Control (D0) | VALO Control (D0) | 3.1527 | 0.0758 | 0.1137 |
| | RIR Control (D0) | RIR Infected (D3) | 1.32 | 0.2506 | 0.2685 |
| | VALO Control (D0) | VALO Infected (D3) | 3.1527 | 0.0758 | 0.1137 |
| | RIR Infected (D3) | VALO Infected (D3) | 1.32 | 0.2506 | 0.2685 |
| | RIR Infected (D3) | RIR Infected (D14) | 6.1929 | 0.0128 | 0.0321 |
| | VALO Infected (D3) | VALO Infected (D14) | 7.1802 | 0.0074 | 0.0321 |
| | RIR Infected (D14) | VALO Infected (D14) | 1.6205 | 0.203 | 0.2538 |
| Shannon Index | RIR Control (D0) | VALO Infected (D0) | 0.0982 | 0.754 | 0.754 |
| | RIR Control (D0) | RIR Infected (D3) | 2.4545 | 0.1172 | 0.1598 |
| | VALO Control (D0) | VALO Infected (D3) | 0.5345 | 0.4647 | 0.5362 |
| | RIR Infected (D3) | VALO Infected (D3) | 0.8836 | 0.3472 | 0.434 |
| | RIR Infected (D3) | RIR Infected (D14) | 2.5929 | 0.1073 | 0.1598 |
| | VALO Infected (D3) | VALO Infected (D14) | 8.0769 | 0.0045 | 0.018 |
| | RIR Infected (D14) | VALO Infected (D14) | 5.9063 | 0.0151 | 0.0283 |

*OTUs, the number of bacteria taxa

Supplementary Table 4. A multivariate ANOVA (PERMANOVA) analysis of β diversity of chicken colon microbiome across experiment groups. To evaluate β diversity the unweighted UniFrac distances were used for PERMANOVA. In Experiment 1, specific pathogen free (SPF) Rhode Island Red (RIR) chickens were used to assess the effect on H9N2 infection on RIR host colon microbiota. Samples were taken at day 0, pre-challenge (D0), day 4 post-challenge (D4), day 10 post-challenge (D10) of experiment In Experiment 2, SPF RIR were compared to SPF VALO chickens to assess differences in host microbiome following H9N2 infection in the two different chicken breeds. Samples were collected at day 0 (D0), pre-challenge, day 3 post-challenge (D3) and day 14 post-challenge(D14) from infected groups. In addition, colon samples were taken in 7 days interval, at day 7 of age (D7), day 14 of age (D14), etc. from RIR and VALO control birds. In both *iv vivo* experiments chickens were challenge with recombinant A/chicken/Pakistan/UDL01/08 H9N2 LPAIV.

| Group 1 | Group 2 | pseudo-F | p-value | q-value |
|---|---|---|---|---|
| Experiment 1 RIR chickens | | | | |
| RIR Control (D0) | RIR Control (D4) | 2.804828 | 0.048 | 0.053053 |
| | RIR Control (D10) | 6.005514 | 0.007 | 0.0182 |
| | RIR Infected (D2) | 2.254156 | 0.046 | 0.053053 |
| RIR Control (D2) | RIR Infected (D2) | 2.555571 | 0.012 | 0.0182 |
| RIR Control (D4) | RIR Infected (D4) | 4.825933 | 0.007 | 0.0182 |
| RIR Control (D4) | RIR Control (D10) | 6.243836 | 0.013 | 0.0182 |
| RIR Control (D10) | RIR Infected (D10) | 5.345121 | 0.010 | 0.0182 |
| Experiment 2. RIR or VALO H9N2-infected chickens versus its controls | | | | |
| RIR Infected (D3) | RIR Control (D3) | 2.560797 | 0.044 | 0.046667 |
| VALO Infected (D3) | VALO control (D3) | 2.210124 | 0.021 | 0.028 |
| VALO Infected (D14) | VALO Control (D14) | 3.991839 | 0.004 | 0.010182 |
| RIR Infected (D14) | RIR Control (D14) | 7.69303 | 0.003 | 0.009333 |
| Experiment 2. RIR-H9N2 infected versus VALO-H9N2 infected chickens | | | | |
| RIR Infected (D0) | VALO Infected (D0) | 2.555234 | 0.017 | 0.018214 |
| RIR Infected (D0) | RIR Infected (D3) | 2.546458 | 0.043 | 0.043 |
| VALO Infected (D0) | VALO Infected (D3) | 4.95074 | 0.009 | 0.012273 |
| RIR Infected (D3) | VALO Infected (D3) | 2.630199 | 0.014 | 0.016154 |
| RIR Infected (D3) | RIR Infected (D14) | 7.099032 | 0.002 | 0.005 |
| VALO Infected (D3) | VALO Infected (D14) | 7.209013 | 0.003 | 0.006429 |
| RIR Infected (D14) | VALO Infected (D14) | 2.780883 | 0.008 | 0.012 |

| | | | | |
|---|---|---|---|---|
| Experiment 2. RIR controls and VALO controls | | | | |
| RIR Control (D7) | RIR Control (D14) | 2.561913 | 0.008 | 0.023333 |
| RIR Control (D7) | RIR Control (D21) | 3.570933 | 0.012 | 0.023333 |
| RIR Control (D7) | RIR Control (D28) | 7.285158 | 0.007 | 0.023333 |
| RIR Control (D7) | RIR Control (D32) | 7.838386 | 0.009 | 0.023333 |
| RIR Control (D14) | RIR Control (D21) | 2.231249 | 0.018 | 0.027 |
| RIR Control (D14) | RIR Control (D28) | 5.935683 | 0.007 | 0.023333 |
| RIR Control (D14) | RIR Control (D32) | 6.453328 | 0.013 | 0.023333 |
| RIR Control (D21) | RIR Control (D28) | 4.368246 | 0.01 | 0.023333 |
| RIR Control (D21) | RIR Control (D32) | 4.554562 | 0.006 | 0.023333 |
| VALO Control (D7) | VALO Control (D14) | 2.281393 | 0.018 | 0.027 |
| VALO Control (D7) | VALO Control (D21) | 2.816923 | 0.011 | 0.023333 |
| VALO Control (D7) | VALO Control (D28) | 4.019477 | 0.011 | 0.023333 |
| VALO Control (D7) | VALO Control (D32) | 4.199853 | 0.011 | 0.023333 |
| VALO Control (D14) | VALO Control (D21) | 2.231236 | 0.022 | 0.030937 |
| VALO Control (D14) | VALO Control (D28) | 2.550725 | 0.024 | 0.032727 |
| VALO Control (D14) | VALO Control (D32) | 2.889433 | 0.014 | 0.023333 |
| VALO Control (D21) | VALO Control (D32) | 2.096668 | 0.025 | 0.033088 |
| Experiment 2. RIR controls versus VALO controls | | | | |
| RIR Control (D7) | VALO Control (D7) | 1.005111 | 0.422 | 0.431591 |
| RIR Control (D28) | VALO Control (D28) | 1.680646 | 0.109 | 0.122625 |
| RIR Control (D21) | VALO Control (D21) | 1.790323 | 0.03 | 0.036486 |
| RIR Control (D32) | VALO Control (D32) | 1.510827 | 0.091 | 0.107763 |

Supplementary Table 5 (excel file). A detailed statistic of analysis of composition of microbiomes (ANCOM) percentile of different bacteria taxa found in Rhode Island Red (RIR) chickens colon across the experimental group. Colon samples were taken at day 0, pre-challenge (D0), day 4 post-challenge (D4) and day 10 post-challenge (D10). RIR chickens were challenge with recombinant A/chicken/Pakistan/UDL01/08 H9N2 LPAIV.
